# Automated Model-Predictive Design of Synthetic Promoters to Control Transcriptional Profiles in Bacteria

**DOI:** 10.1101/2021.09.01.458561

**Authors:** Travis La Fleur, Ayaan Hossain, Howard M. Salis

## Abstract

Transcription rates are regulated by the interactions between RNA polymerase, sigma factor, and promoter DNA sequences in bacteria. However, it remains unclear how non-canonical sequence motifs collectively control transcription rates. Here, we combined massively parallel assays, biophysics, and machine learning to develop a 346-parameter model that predicts site-specific transcription initiation rates for any σ^70^ promoter sequence, validated across 17396 bacterial promoters with diverse sequences. We applied the model to predict genetic context effects, design σ^70^ promoters with desired transcription rates, and identify undesired promoters inside engineered genetic systems. The model provides a biophysical basis for understanding gene regulation in natural genetic systems and precise transcriptional control for engineering synthetic genetic systems.

**One-Sentence Summary:** A 346-parameter model predicted DNA’s interactions with RNA polymerase initiation complex, enabling accurate transcription rate predictions and automated promoter design in bacterial genetic systems.

## Main Text

Transcription is the gene expression process responsible for producing all RNA and is a common engineering target for creating novel products, including microbial chemical factories, toxin-sensing genetic circuits, and mRNA vaccines ^1-3^. However, while DNA assembly techniques enable the construction of custom-designed genetic systems ^4^, it remains challenging to *a priori* predict and control a system’s gene expression profile ^5^, for example, by initiating transcription with desired rates at specific DNA start sites, while minimizing transcription from all other DNA sequence regions. Currently, transcriptional control relies on empirical characterization of promoters as modular genetic parts ^6^. Applying modular design to transcriptional control ignores other sources of transcription, for example, inside coding regions, as well as local and long-distance interactions that alter transcription rates and start sites in unexpected ways ^7^. Poorly controlled transcriptional profiles can lead to malfunctioning genetic systems, including the undesired production of anti-sense RNA and truncated proteins as well as the misbalancing of protein expression levels that lead to lower system activities ^8, 9^.

A key challenge is to quantitatively predict how polymerase initiation complex – RNA polymerase (RNAP) and a sigma factor (σ) in bacteria – interacts with arbitrary DNA sequences ^10, 11^. Beyond canonical sequence motifs, it remains unclear how the strengths of multiple interactions collectively determine transcription initiation rates and start sites, particularly when bound to RNAP/σ^70^, which initiates transcription at the majority of bacterial promoters. To help resolve this challenge, we carried out massively parallel experiments on designed promoter sequences to systematically measure the interactions controlling site-specific transcription at σ^70^ promoters (**Figure 1A**). With this data, we developed a statistical thermodynamic model that calculates how RNAP/σ^70^ interacts with arbitrary DNA to predict transcription initiation rates at each position. The model has only 346 interaction energy parameters, but accurately predicts the transcription rates of 17396 bacterial promoters with diverse sequences. We show how the model enables the automated design and debugging of transcriptional profiles in engineered genetic systems.

**Figure 1:**
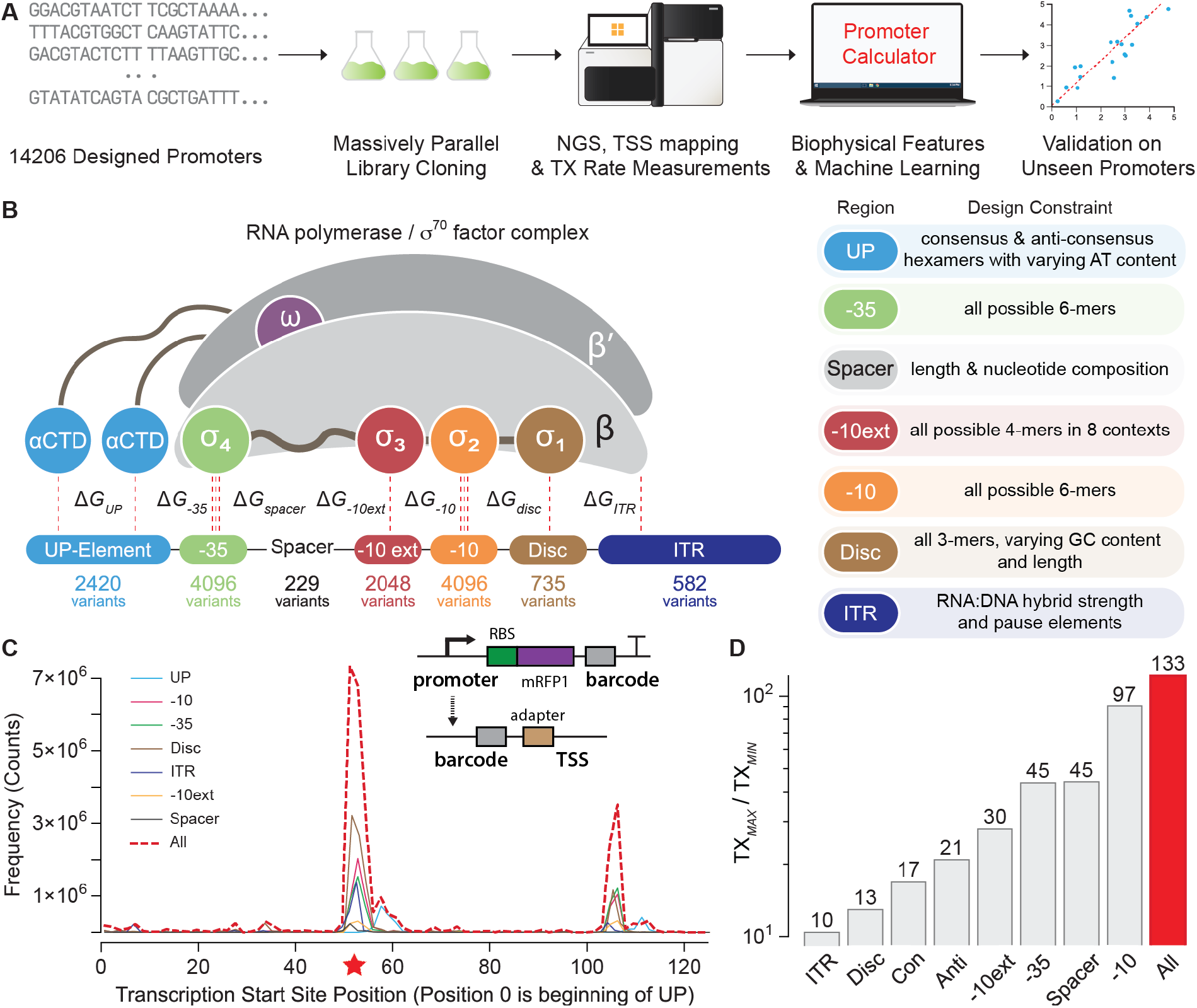
Massively Parallel Transcription Rate and Start Site Measurements. (**A**) Model development combined promoter design, barcoded oligopool synthesis, library cloning & culturing, and next-generation sequencing to measure the transcription start site and transcription rate of each promoter variant. (**B**) The interaction strengths between RNAP/σ^70^ and promoter DNA control transcription initiation rates. 14206 promoter variants were designed to quantify how sequence modifications affect each interaction. Sequence design criteria are shown. (**C**) The frequencies of observed transcription start sites are shown for each set of promoter variants. The star indicates the predominant start site. The inset schematic shows the system architecture, the locations of the promoter and barcode variants, and the cDNA architecture after library preparation. (**D**) The transcription rates’ dynamic ranges are shown for each set of promoter variants, considering only the predominant start site. Con: UP-varied promoters with consensus hexamers. Anti: UP-varied promoters with anti-consensus hexamers.

To begin, we designed 14206 promoter variants with varied motif sequences to systematically perturb the interactions that affect RNAP/σ^70^ binding and transcriptional initiation (**Figure 1B**). These interactions occur at DNA sites known by their canonical positions ^12, 13^, including (i) an upstream 6-nucleotide site called the -35 motif; (ii) a 20-nucleotide region that appears upstream of the -35 motif, called the UP element; (iii) a downstream 6-nucleotide site called the -10 motif; (iv) a spacer region that separates the -10 and -35 motifs; (v) a 4-nucleotide site upstream of the - 10 motif, called the -10 extended motif (−10ext); (vi) a typically 6-nucleotide region in between the -10 motif and transcriptional start site, called the discriminator (Disc); and (vii) the first 20 transcribed nucleotides, called the initial transcribed region (ITR).

Initial RNAP/σ^70^ binding to a promoter is controlled by the interaction Gibbs free energies (ΔG) at the UP, -35, -10 extended, and -10 motifs as well as the torsional stress controlled by the length of the spacer region ^12-17^. Bound RNAP/σ^70^ then undergoes a conformational change that catalyzes double-stranded DNA separation, creating a transcription bubble that initially encompasses half the -10 motif, the Disc, and the first two nucleotides of the ITR ^13, 14^. RNA polymerization begins at a transcription start site (TSS) determined by where the catalytic site in the β subunit contacts the DNA template, canonically at position +1. The transcription bubble is then stabilized by interactions in the Disc ^18^ and the formation of an R-loop, whereby the newly synthesized RNA strand immediately hybridizes to the DNA template ^19^. The ITR sequence controls the R-loop’s thermodynamic stability. Finally, transcription initiation is successful once enough DNA is pulled into the stable transcription bubble that the accumulated stress exceeds the interaction strength between RNAP/σ^70^ and promoter DNA, causing promoter escape and a transition to processive RNA synthesis ^19-21^.

From these interactions, we formulated a statistical thermodynamic model of transcriptional initiation that accounts for competitive binding of RNAP/σ^70^ to all DNA, the multiple sequence contacts that control RNAP/σ^70^ recruitment at each promoter, and the multiple internal states during transcription initiation (**Supplementary Information**). So long as the internal states do not become abundant, for example, by significant transcriptional pausing ^22^, the model indicates that we can decompose how a promoter’s sequence controls the interaction energies into a sum of free energies that can be related to the transcription initiation rate (*TX*), according to:

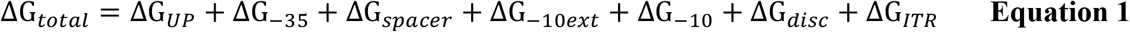

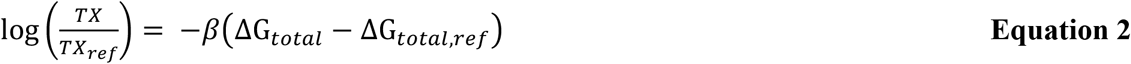

where ΔG_*total*_ is the difference in free energy between an unbound promoter and a promoter-RNAP/σ^70^ complex with a stable transcriptional bubble, which is used to predict a promoter’s TX rate in comparison to a reference promoter sequence with calculated ΔG_*total,ref*_ and measured *TX*_*ref*_. β is a measurable model constant that converts free energies into state probabilities. Here, we arbitrarily set ΔG_*total,ref*_ to zero so that stronger (weaker) interaction free energies have more negative (positive) values in comparison.

We designed the 14206 promoter sequences to measure how each motif sequence alters these free energies to create a sequence-complete model, including all possible -10 hexamers, all possible - 35 hexamers, all possible -10 extended motifs within 8 consensus and anti-consensus background configurations, 229 sequence spacers with varied lengths and nucleotide compositions, 2420 UP elements with varied AT content within 4 consensus and anti-consensus background configurations, 735 discriminator elements with varied lengths and GC content, and 582 ITR sequences with varied R-loop stabilities (**Figure 1B**). Using oligopool synthesis and two-step library cloning, we constructed a barcoded plasmid pool that uses each promoter in a common genetic context, expressing a single protein with a moderate translation initiation rate (about 5000 on the RBS Calculator v2.1 scale ^5^), with unique barcode sequences positioned in the 3’ untranslated region to avoid confounding effects.

We then carried out *in vitro* transcription reactions, TSS mapping, and next-generation sequencing to measure the TX rate of each promoter at each TSS, utilizing a minimal system containing only RNAP/σ^70^, the barcoded plasmid pool, NTPs, and buffer (**Methods**). TSS mapping was performed by harvesting product RNA, ligating a 5’ RNA adapter, converting to cDNA, circularizing, using PCR to generate amplicons containing the barcode and TSS, and obtaining over 323 million barcode-TSS mapped reads from Illumina sequencing. Across all promoters, we found 182 TSSs with at least 1000 mapped reads, revealing a primary TSS region and a less frequent, secondary TSS region arising from a downstream cryptic promoter. TX rates were then measured by carrying out DNA-Seq and RNA-Seq on triplicate *in vitro* transcription reactions, separately harvesting DNA and RNA, converting RNA to cDNA, using PCR to generate barcode-containing amplicons, and obtaining between 81.3 and 130.5 million barcoded reads per reaction. From the RNA/DNA read count ratios, we obtained TX rates for 13480 promoter variants (mean and standard deviation), excluding any with fewer than 50 read counts in any replicate. Replicate read count measurements were highly reproducible (R^2^ = 0.89 to 0.99, **Figure S1**) with 5388 high-precision TX rates (coefficient of variation < 0.40). All promoter sequences, DNA read counts, RNA read counts, TX rates, and TSS frequencies are found in the **Supplementary Data**.

We began model training by identifying 5193 promoter variants where RNAP/σ^70^ bound to a single site with one predominant TSS, enabling us to unambiguously pinpoint their motif sequences (**Figure 1C, Figure S2**). The TX rates for these single-site promoters altogether varied by 133-fold with sizable effects from each individual motif (**Figure 1D**). Importantly, endogenous RNAses were not present in the *in vitro* transcription reactions, enabling us to vary the Disc and ITR sequences without changing the mRNA’s stability, which is not possible in equivalent *in vivo* measurements. We then specified 472 sequence, structural, and energetic properties to relate how each motif sequence contributes to each binding free energy term (**Table S1**). For example, for any UP sequence, we calculate the minor groove width of the distal and proximal UP sites ^23^; for any ITR sequence, we calculate the thermodynamic stability of the R-loop ^24, 25^; for any spacer sequence, we calculate the local DNA rigidity ^26^. As categorical properties, we split the -35, -10, and Disc motifs into six 3-nucleotide regions and include 384 3-mers. We split the -10ext motif into 2-nucleotide regions and include 32 2-mers. We also include the spacer length as a categorical property.

We then randomly split our dataset into a training set (4673 promoters, 90%), carried out 10-fold cross-validation to identify the optimal hyperparameters of a machine learning model, and tested its accuracy on the remaining unseen test set (520 promoters, 10%). We evaluated several models (**Table S2**) that use regularization to identify the optimal coefficients of a linear, additive model that mirrors our free energy parameterization (**Equation 1**) with dataset normalization that mirrors our statistical thermodynamic model (**Equation 2**). We carried out feature reduction to remove extraneous properties, which lowered the number of fitted coefficients. The pruned properties included the first 2-nucleotide region of the -10ext motif and the last 3-nucleotides of the Disc motif, which had no discernable effect on TX rate in this dataset. Overall, we found that a ridge regression model with 346 fitted coefficients yielded a convergent learning curve (**Figure 2A**) and highly accurate predictions (*R*^2^ = 0.80, **Figure 2B**) with similarly low error distributions (**Figure 2C**) across both the training and unseen test datasets. We then performed ANOVA analysis to quantify how each predicted free energy term contributed to the promoters’ measured TX rates in our dataset and found that 83% of the TX rate variance could be explained by varying the interactions that control initial RNAP/σ^70^ recruitment, using the UP, -35, -10, -10ext, and spacer motifs (**Figure 2D**). In contrast, 7.1% of the TX rate variance was explained by differences in the interactions controlling DNA melting, R-loop formation, and promoter escape, which are affected by the Disc and ITR regions.

**Figure 2:**
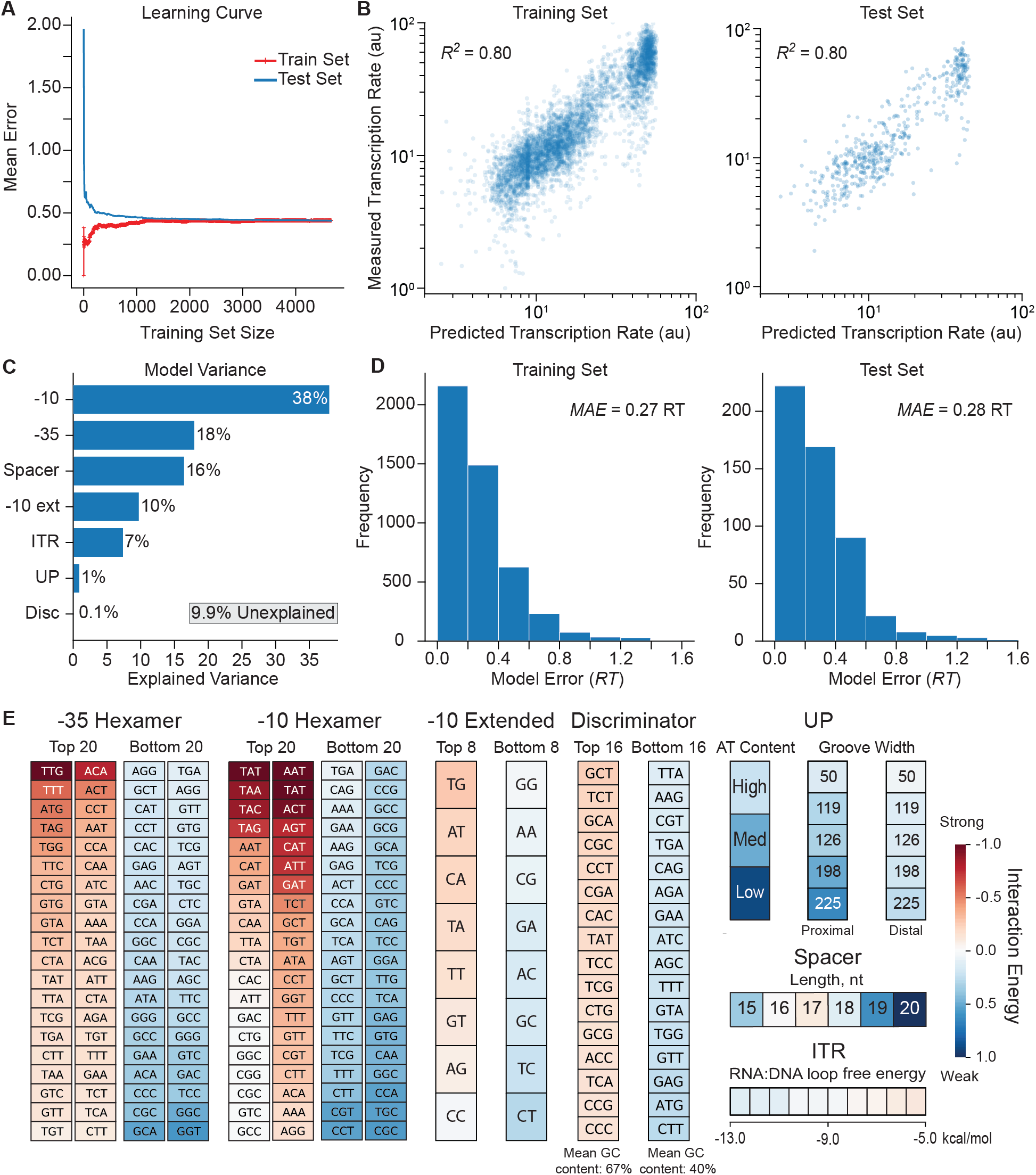
Model Development and Validation using Machine Learning. (**A**) A learning curve shows the training and testing of a ridge regression model to identify the unknown interaction energies. (**B**) Model-predicted transcription rates are compared to measured transcription rates for both (left) the training set and (right) the test set. (**C**) The explained variances for each promoter interaction are shown. (**D**) Histograms of model error are shown for the training and test sets, using energy units. MAE is mean absolute error. (**E**) The learned interaction energies are shown for the strongest and weakest ones. A poster-sized schematic showing all interaction energies is available (**Supplementary Information)**.

A key benefit of our hybrid biophysical-machine learning approach is that the fitted coefficients quantify the physical interactions between RNAP/σ^70^ and each promoter motif (**Figure 2E**), including all motif sequences and enabling direct comparisons to already known canonical interactions as positive controls. For example, without human supervision, our approach correctly identified the canonical -35 motif (TTGACA), -10 motif (TATAAT), extended -10 motif (TG), and the optimal spacer length (17 base pairs). We also found that AT-rich distal and proximal UP sites had more favorable interactions and that GC-rich Disc motifs were enriched, which were previously observed ^17, 18^. However, most promoters do not contain canonical motif sequences; their TX rates are controlled by a mixture of weaker interactions. A key novelty of our model is its complete set of interaction energies, covering all possible promoter sequences, which provides the ability to predict the TX rate of any σ^70^ promoter. All interaction energies are available in **Supplementary Data**, including numerical values and a poster-sized chart.

We then tested the model’s generalizability and accuracy across 17396 well-characterized promoters, using sequence information alone to predict their transcriptional profiles, specifically their TX rates for each potential TSS position (**Figure 3A**). Here, we do not input the promoters’ actual TSS positions and motifs, and instead combine the interaction energies with statistical thermodynamics to identify the most likely RNAP/σ^70^ binding site configuration. For each potential TSS position, we scan the surrounding DNA sequence and enumerate the several ways that RNAP/σ^70^ may bind to it, varying the spacer and Disc lengths with corresponding changes in motif sequences. We apply the model’s interaction energies to calculate the ΔG_total_ for each configuration (**Equation 1**) and determine the configuration with the most negative ΔG_total_, which is then used to predict the TX rate (**Equation 2**). We repeat these calculations for each potential TSS position within the inputted DNA sequence.

**Figure 3:**
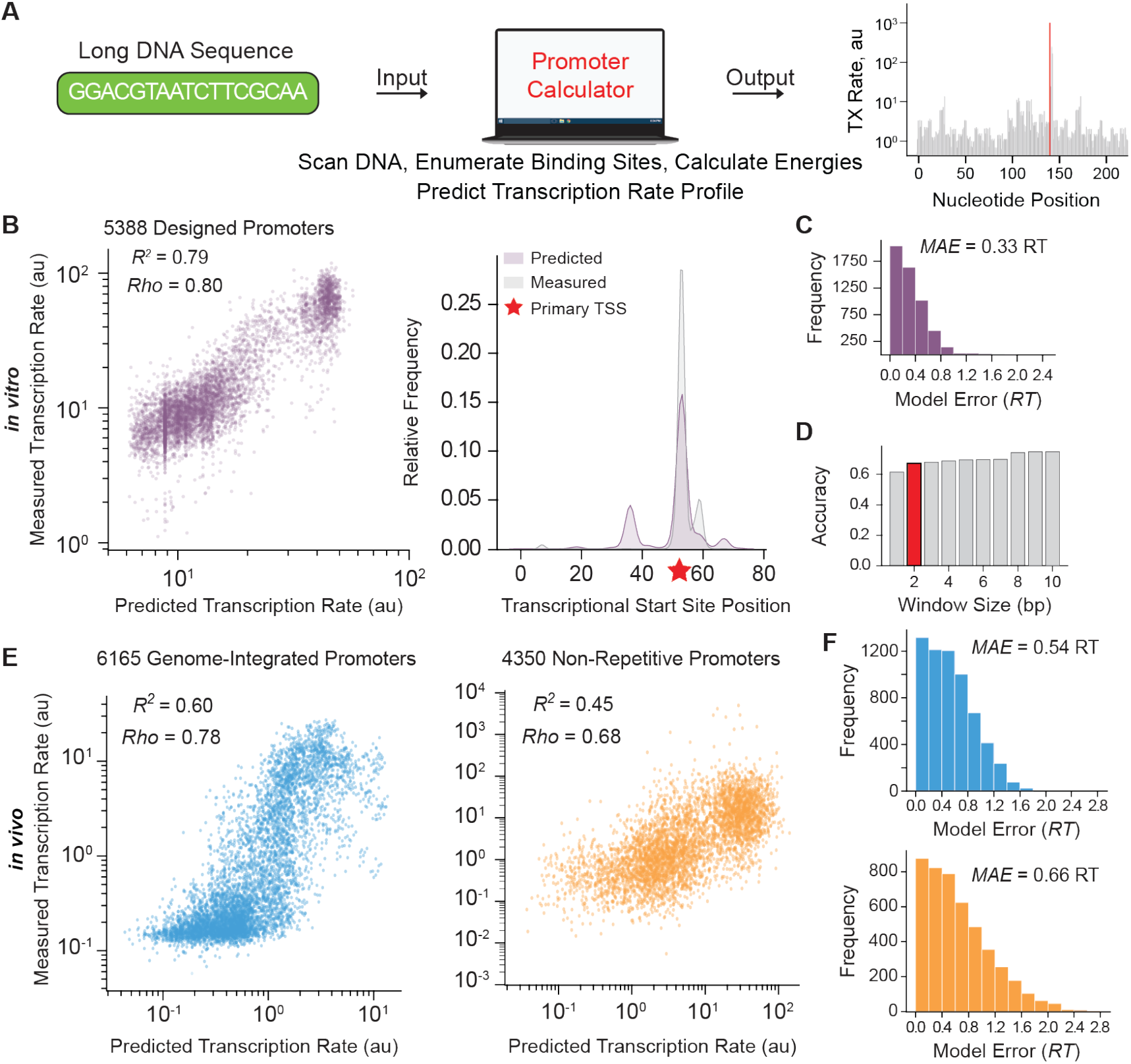
Validation of Sequence-to-Function Model on Diverse Promoters. (**A**) Arbitrary DNA sequences are inputted into the model to predict its transcriptional profile (transcription rates vs. nucleotide positions) without start site information. (**B**) Model predictions for 5388 designed promoters (this study) are compared to *in vitro* transcription rate and start site measurements. (**C**) The histogram of the scanning model’s error is shown when evaluated on the designed promoters. (**D**) The model’s start site prediction accuracy is shown within defined windows. (**E**) Model predictions on unseen test sets, including (left) 4350 non-repetitive promoters ^27^ and (right) 6185 genome-integrated modular promoters ^28^, are compared to *in vivo* transcription rate measurements. (**F**) Histograms of the model’s error when evaluated on (top) 4350 non-repetitive promoters and (bottom) 6185 genome-integrated modular promoters.

We carried out these additional tests on four datasets: (1) 5388 designed promoters with high-precision TX rate and TSS measurements, characterized here using *in vitro* RNAP/σ^70^ transcription assays; (2) 6165 promoters using combinations of motifs to express a ribozyme-insulated transcript, characterized using genome-integrated test circuits inside *E. coli* cells ^28^; (3) 4350 non-repetitive promoters using highly diverse sequences to express a transcript, characterized using test circuits carried on multi-copy plasmids inside *E. coli* cells ^27^; and (4) 1493 inducible promoters that contain transcription factor binding sites and express a ribozyme-insulated transcript, also characterized using genome-integrated test circuits inside *E. coli* cells ^29^. We first confirmed that the statistical thermodynamic model accurately predicted the TX rates of the *in vitro* dataset #1 without utilizing TSS measurements (R^2^ = 0.79, ρ = 0.80, **Figure 3B**) with 4954 promoters (92%) having predicted TX rates to within 2-fold of their actual TX rates (**Figure 3C**) and TSS positions predicted to within 2 bp (67% accuracy, **Figure 3D**). Next, we tested the model’s predictions on the three *in vivo* datasets, which were not used during model development. We found that the statistical thermodynamic model retained accuracy (R^2^ = 0.60, ρ = 0.78; R^2^ = 0.45, ρ = 0.68; R^2^ = 0.65, ρ = 0.69; **Figure 3E** and **Figure S3**) with low model errors (**Figure 3F**), even though *in vivo* mRNA levels are affected by other potentially confounding factors, for example, variation in DNA copy number and mRNA stability.

To evaluate the model’s predictions using additional types of measurements, we then selected ten of the non-repetitive promoters in dataset #3 and characterized their expression levels in test genetic circuits inside *E. coli* cells, utilizing RT-qPCR to measure their mRNA levels and flow cytometry to measure their fluorescent reporter protein levels. We found that the model’s predicted TX rates for these promoters were highly proportional to their measured mRNA levels (R^2^ = 0.75, ρ = 0.87) and fluorescent protein expression levels (R^2^ = 0.78, ρ = 0.87) over a 96000-fold dynamic range (**Figure S4**). All dataset sequences, model predictions, and TX rate measurements are included in **Supplementary Data**.

Across microbial biotechnology, promoters are routinely treated as reusable, swappable genetic parts to control protein expression levels ^6^. However, the model predicts that the DNA sequences surrounding a promoter will greatly affect its TX rate, specifically the UP and ITR regions, which would explain why some promoters express certain proteins at much lower levels. We next tested the model’s ability to predict, explain, and overcome these genetic context effects. First, we designed UP and ITR sequences that the model predicted would greatly alter a promoter’s TX rate, inserted them around the commonly used J23101 promoter, and measured their *in vivo* TX rates using a fluorescent protein reporter inside *E. coli* cells (**Methods**). Changing only the UP region or only the ITR region reduced the promoter’s TX rate by up to 5.3-fold, while changing both regions reduced the promoter’s TX rate by up to 9.4-fold (**Figure 4A**). The model predicted that these modifications led to the emergence of off-target TSSs that may interfere with transcriptional initiation ^9^ (**Figure 4B**). We next evaluated whether these genetic context effects are potentially widespread by carrying out Monte Carlo simulations to predict the range of TX rates expected when varying the UP and ITR regions of several commonly used promoters (**Figure 4C**). Promoters with higher average TX rates exhibited much higher sensitivity to changing the UP and ITR regions with an overall average coefficient of variation of 0.40. We then tested whether the model is able to correctly account for genetic context effects and predict the promoters’ TX rates in a particular context. To do this, we selected a classic community dataset whereby the *in vivo* TX rates of these commonly used promoters were previously characterized inside *E. coli* cells and within a specific genetic context ^4^. We then inputted the specific UP, core promoter, and ITR sequences from this dataset into our model and confirmed accurate TX rate predictions (*R*^2^ = 0.72, ρ = 0.86, **Figure 4D**), further illustrating the model’s generality.

**Figure 4:**
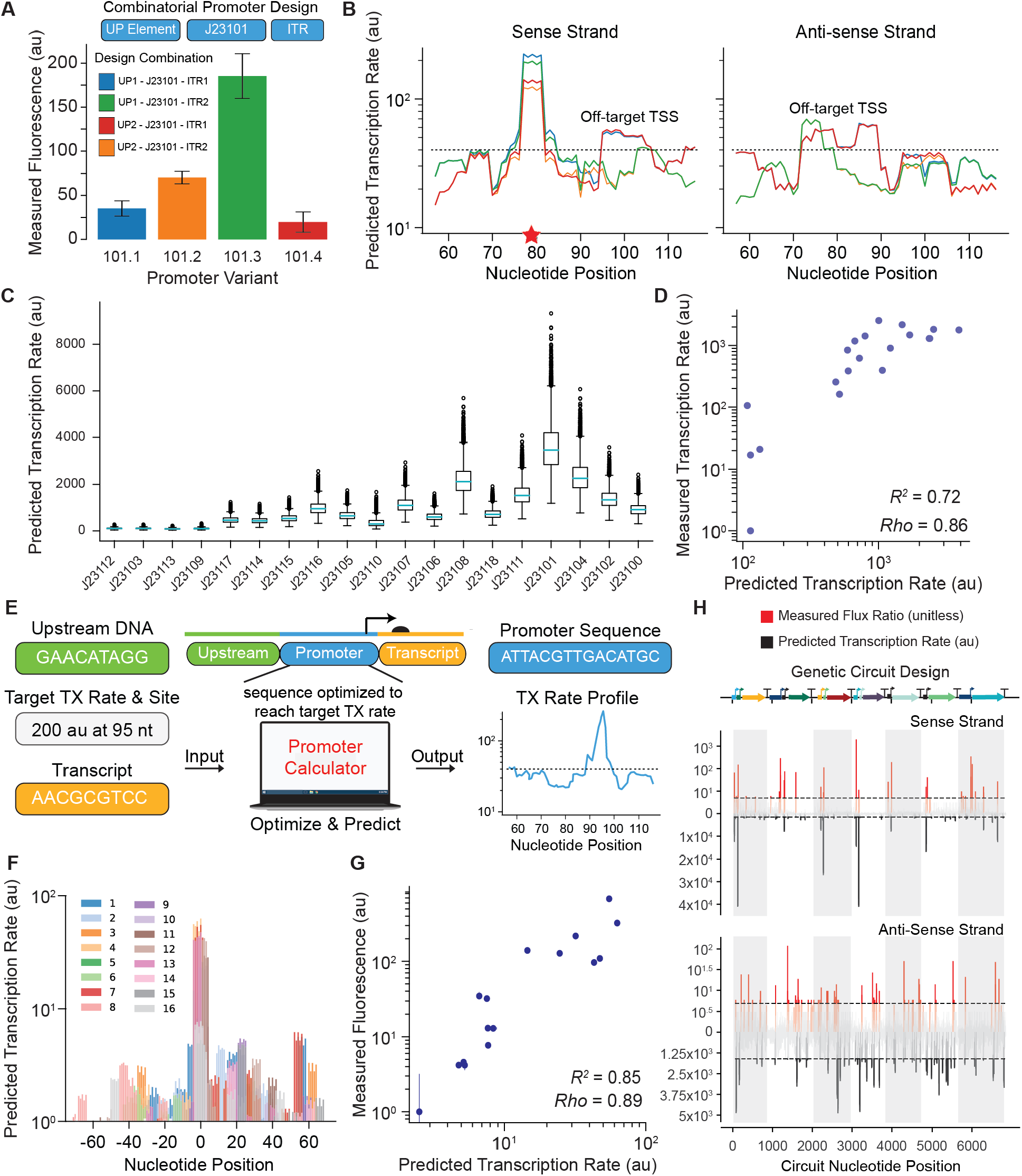
Overcoming Context Effects to Design and Debug Transcriptional Profiles. (**A**) Measured *in vivo* transcription rates for the J23101 promoter genetic part with varied upstream (UP) and downstream (ITR) sequences, showing transcriptional context effects. (**B**) Transcription rate profiles are predicted for each J23101 promoter variant, showing predominant and off-target transcription start sites for the (left) sense and (right) anti-sense strands. The star shows the expected start site. (**C**) Transcriptional context effects are quantified for common promoter genetic parts, showing predicted transcription rates when varying upstream and downstream sequences. (**D**) Model predictions are compared to *in vivo* transcription rate measurements for these promoters when specifying their upstream and downstream sequences. (**E**) Promoter sequences are automatically designed with desired transcription rates and start sites, including the upstream and downstream sequences (**F**) Predicted transcriptional profiles for 16 automatically designed promoters with varied transcription rate specifications. (**G**) Model predictions are compared to *in vivo* transcription rate measurements for the 16 designed promoters. Dots and bars are the mean and standard deviation from duplicate measurements. (**H**) Predicted σ^70^ transcriptional profiles of an 11-promoter genetic circuit ^8^ are compared to *in vivo* transcription rate and start site measurements on the (top) sense and (bottom) anti-sense strand. Transcription flux ratios are the measured differences in adjacent mRNA levels from RNA-Seq. White and grey shadows correspond to each transcribed cistron. The dotted black line shows the minimum transcription rates that define a start site.

Next, we developed an automated optimization algorithm that designs promoter sequences with desired transcriptional profiles, taking into account both the upstream DNA sequence and the transcribed mRNA sequence (**Figure 4E**). The algorithm identifies optimal promoter sequences that provide user-defined TX rates at desired TSSs, while minimizing the TX rate at undesired, off-target TSSs (**Methods**). We tested this algorithm by designing, constructing, and characterizing 16 synthetic promoter sequences with targeted, systematically varied TX rates. The designed promoters were 77 base pairs long and shared only 26% sequence similarity (average pairwise hamming distance of 57). The model predicted single predominant TSSs at the desired locations, though it becomes progressively more difficult to achieve single-TSS predominance as the targeted TX rate is lowered (**Figure 4F**). We then measured the promoters’ TX rates, each expressing a ribozyme-insulated transcript on a multi-copy plasmid inside *E. coli* cells during exponential growth (**Methods**), and found that the model’s predictions were highly accurate (R^2^ = 0.85, ρ = 0.89, **Figure 4G**), enabling fine control of site-specific transcription rates over a 682-fold range, while taking into account the surrounding genetic context.

Finally, we demonstrated how the statistical thermodynamic model facilitates the debugging of large genetic circuits by predicting their transcriptional profiles and identifying cryptic promoters that could disrupt the circuit’s function. As a demonstration, we selected a recently characterized genetic circuit that uses 11 engineered promoters and 7 transcription factors to carry out digital logic ^8^ (**Figure 4H**). The model predicted single-TSS peaks for 9 of the engineered promoters and identified 51 cryptic promoters (18 sense and 33 anti-sense). These predictions were qualitatively confirmed by system-wide RNA-Seq measurements and transcriptional flux inferences with an overall accuracy of 55% for cryptic promoter identification (**Methods**). Future design software can leverage these predictions to redesign protein coding regions and remove cryptic promoters before genetic circuits are constructed.

In conclusion, we created a parsimonious (human-understandable) biophysical model of bacterial transcription initiation with demonstrated accuracy across thousands of σ^70^ promoters with highly diverse sequences, enabling site-specific control over TX rates in engineered genetic systems. Our model shows how multiple weak interactions contribute to RNAP/σ^70^ transcriptional control, including its promiscuous activity on DNA sequences without cognate binding motifs ^30^. Our bottom-up model-building approach is readily extendable to other RNAP/σ complexes and demonstrates how advances in machine learning can enhance, rather than replace, existing thermodynamic formalisms with the overall goal of creating a universal system-wide language for engineering gene regulation in synthetic genetic systems.

## Online Methods

### Library Design

14206 σ^70^ promoters were rationally designed to include 2420 UP elements, 4096 -35 hexamers, 229 spacers, 2048 -10 extended motifs, 4096 -10 hexamers, 735 discriminators, and 582 initial transcribed regions (ITR). The 2420 UP element sequences were designed in two subsets: (i) the first set varied adenine and thymine (AT) content from 0-100% within a background sequence containing combinations of consensus, anti-consensus, and mutated -10 and -35 hexamers; and (ii) the second set introduced short cytosine and adenine repeats into the distal and proximal binding sites, while varying AT content within a background sequence containing the same combinations of consensus, anti-consensus, and mutated hexamers. The -10 hexamer set was designed to include the 4096 6-nt sequences in the -10 hexamer, a consensus -35 hexamer, and a 17-bp spacer. The - 35 hexamer set was designed to include the 4096 possible 6-nt sequences in the -35 hexamer, a consensus -10 hexamer, and 17-bp spacer. The 229 spacers were designed by varying the spacer length from 1-to 32-bp, in between a consensus -35 and a consensus -10 hexamer, while varying the spacer nucleotide composition at each length. The -10 extended promoter variants were designed to include the 256 possible 4-nt sequence at positions -14 to -17 within 8 different background sequences containing combinations of consensus, anti-consensus, and mutated -10 and -35 hexamer sequences. The 735 discriminator variants were designed in two sets with a background sequence containing a consensus -10 hexamer: (i) a first set that varied the guanine-cytosine (GC) content from 0-100% and the discriminator length from 6 to 8 bp with 5 randomly generated sequences satisfying each criteria; and (ii) a second set that introduced all possible 3-nt sequences in the first half of the discriminator and varied the GC content in the remaining 4-nt. The 582 ITR variants were designed to vary the number of purine bases, the GC content, and the presence of -10 hexamer-like pause sequences within the ITR. Unless otherwise stated, promoter variants were designed within a background containing consensus hexamers and a canonical 17-bp spacer. A complete list of the promoter sequences is available in the **Supplementary Data**.

### Oligo Pool Design and Optimization

An oligopool containing a mixture of 170-nt oligonucleotides was synthesized (Genscript). Each oligonucleotide design contained a promoter variant, four restriction enzyme cut sites, two primer binding sites, and a unique 20 nucleotide barcode sequence (**Figure S5A**). A custom design algorithm was used the create barcodes and primer binding sites. Barcodes were designed to maximize pair-wise hamming distance between each other and with respect to the unique *k*-mer list generated from the 14206 designed promoters and the *E. coli* genome. Primer binding site sequences were designed to have target melting temperatures of 55°C, low probabilities of forming primer dimers, and minimal primer structure. Primer binding site sequences were also designed with maximum dissimilarity to barcode sequences, promoter sequences, and the *E. coli* genome sequence.

### Library Cloning and Growth

PCR was carried out using the oligopool as DNA template to create an insertion cassette library, using Q5 High-Fidelity DNA Polymerase (NEB), 20 cycles, and primers 1 and 2 (**Table S3**). DNA was extracted and purified via gel electrophoresis. The purified insertion cassette library and plasmid pFTV1 were double-digested using BamHI and HindIII (NEB), purified, ligated, purified again, and then transformed into NEB 5-alpha Electrocompetent *E. coli* (NEB). All ligations used T7 DNA ligase (NEB) unless otherwise stated. Transformation recovery was 60 min at 37°C using 1 mL of pre-warmed SOC media. Transformation efficiency was 1.2e10^7^ CFU per mL recovery broth. The transformed cell library was used to inoculate duplicate cultures using 5 mL LB media supplemented with 50 ug/mL chloramphenicol. Shaking cultures were incubated overnight at 37°C. One culture was used for cryostocking. Plasmid purification was performed on the other culture using an EZNA plasmid mini kit (Zymo). The plasmid library was sequentially digested with SpeI and EcoRI, followed by rSAP treatment. Digested plasmid pool was then purified via gel extraction.

To generate DNA templates for carrying out *in vitro* transcription and TSS mapping, the digested plasmid pool was ligated to a SpeI/EcoRI-digested DNA fragment containing a ribosome binding site (RBS) and mRFP1 coding sequence (**Figure S5B, Table S3**), followed by purification and transformation into NEB 5-alpha Electrocompetent *E. coli* cells. The expression plasmid library (**Figure S5C**) was then harvested from overnight shaking cultures grown in LB media supplemented with 50 ug/mL chloramphenicol. For quantification of library coverage, PCR was carried out on the plasmid library to generate promoter-barcode amplicons, followed by next-generation sequencing (Illumina MiSeq), barcode mapping, and promoter counting. 1ug aliquots of the expression plasmid library were used as DNA template for the *in vitro* transcription reactions.

### *in vitro* Transcription Rate Measurements

Replicate *in vitro* transcription reactions were carried out for 3 hours at 37°C, combining 1 ug of the expression plasmid library, 0.5 mM of each NTP, 1X *E. coli* RNA Polymerase Reaction Buffer, and 1 uL of *E. coli* RNA Polymerase Holoenzyme (NEB). DNA was removed by adding 2.5 units of TURBO DNase and incubating at 37°C for 30 min. The RNA product was purified using a RNA Clean & Concentrator Kit (Zymo).

For quantification of RNA variant levels after transcription (RNA-Seq), cDNA first-strand synthesis reactions were first carried out on the harvested RNA, using 200 units of SuperScript IV (Invitrogen), 1X buffer, and transcript-specific primer 2 (**Table S3**). PCR was carried out to generate barcode-containing amplicons (121 bp), using 5 ng of cDNA as template, primers 2 and 3, and 25 cycles of amplification (**Table S3**), followed by next-generation sequencing (Illumina HiSeq 2500, 150 bp paired-end), barcode mapping, and barcode counting. For quantification of DNA variant levels in the expression plasmid library (DNA-Seq), PCR was first carried out on 10 ng of the expression plasmid library to generate promoter-barcode amplicons (847 bp long), using primers 1 and 2 and 25 cycles of amplification. The promoter-barcode regions were then condensed into shorter amplicons (271 bp) by treating the PCR product with T4 Polynucleotide Kinase (NEB), carrying out DNA circularization by ligation using T4 DNA Ligase (NEB), purifying the monomer DNA circles via gel extraction, and then PCR amplifying using primers 3 and 4 with 25 cycles (**Table S3**). The promoter-barcode amplicon library was sequenced using Illumina HiSeq 2500 (150 bp, paired-end), followed by barcode mapping, promoter identification, and counting. Three independent *in vitro* transcription reactions, RNA-Seq, and DNA-Seq assays were carried out.

### Transcriptional Start Site Mapping

Starting with RNA harvested from the *in vitro* transcription reactions, the locations of transcriptional start sites (TSSs) were determined by treating the RNA with TURBO DNase to remove template DNA, purifying, treating with RNA 5’ polyphosphatase (Epicentre) to remove 5’ phosphates, purifying, ligating the RNA to an 5’ RNA adapter (**Table S3**) using T4 RNA Ligase (NEB), and purifying again. The treated RNA product was then converted to cDNA by treating with SuperScript IV (Invitrogen) and the transcript-specific primer 2 (**Table S3**). PCR was then carried out using 10ng of cDNA as template, primers 2 and 5 (**Table S3**), and 30 cycles of amplification. The linear DNA product was then circularized by first treating it with T4 Polynucleotide Kinase, followed by ligating using T4 DNA Ligase. Monomer DNA circles were gel extracted. PCR was then carried out to generate the barcode-TSS amplicon library, using primers 3 and 6 in 25 cycles of amplification. The barcode-TSS amplicon library was then sequenced using Illumina HiSeq 2500 (150 bp, paired-end).

### Analysis of Next-generation Sequencing Data

Barcode mapping, barcode counting, and promoter identification were carried out using custom software developed to analyze data from massively parallel reporter assays. The abundances of the cDNA-derived RNA transcript variants were quantified by counting the numbers of unique barcodes that were within 1 edit distance of the expected barcode sequences. The numbers of each DNA variant within the expression plasmid library were determined by first identifying associated barcode sequences that perfectly matched the expected barcode sequences, extracting the promoter sequence region, and mapping it to the expected promoter sequence according to barcode association. DNA counts were included if the promoter sequence perfectly mapped to the expected sequence. The RNA transcript counts and DNA counts were used to determine the relative transcription initiation rate of each promoter variant *i* in replicate *j* according to the formula:

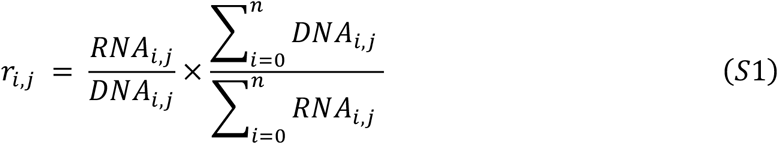

where RNA_i,j_ is the cDNA-derived barcode count for variant *i* in replicate *j*, DNA_i,j_ is the DNA count containing promoter variant *i* in replicate *j*, and *n* is the total number of variants in replicate *j*. Replicate comparisons for RNA barcode counts, DNA promoter counts, and transcription rate measurements are shown in **Figure S1**. Transcriptional start site locations were determined using the TSS-barcode reads. For each read, the barcode was mapped and used to identify the associated reference promoter. The TSS location was then mapped by first identifying the position of a known constant flanking sequence, extracting the adjacent TSS-containing sequence, and carrying out a Smith-Waterman alignment using the associated promoter sequence as a reference. TSS locations were only counted when the TSS-containing sequence contained only one start location that perfectly matched the expected sequence.

### Data Filtering

For model training, the set of promoter variants with a single, predominant transcription start site was defined by the intersection of three criteria: (i) promoter variants must have with at least 50 RNA-Seq and 50 DNA-Seq counts across all triplicate measurements; (ii) promoter variants must have one predominant TSS with at least twice the counts as the next most predominant TSS; and (iii) promoter variants must have a predominant TSS within a 10-bp window surrounding the anticipated TSS location. Data filtering yielded 5193 promoter variants that satisfied these criteria. For model testing, the set of promoter variants with high-precision transcription rate measurements was defined using a single criterion: all promoter variants must have coefficients of variation (CVs) that are less than 0.40. CVs are calculated by determining the standard deviation of the transcription rate measurements and dividing by the mean of the transcription rate measurements across the three independent measurements. Data filtering yielded 5388 promoter variants that satisfied this criterion.

### A Biophysical Model of Bacterial Transcriptional Initiation

We describe the process of prokaryotic transcriptional initiation at a single promoter as a 3-state system (**Figure S6**). In state 0, RNAP/σ is not bound to the promoter DNA (free state). In state 1, RNAP/σ is bound to the promoter DNA, creating a complex in the closed conformational state. In state 2, RNAP/σ is bound to the promoter DNA with a stable transcriptional bubble, which includes the transition from the closed-to-open conformation, the initial melting of promoter DNA to create an unstable transcriptional bubble, and the formation of a stable transcriptional bubble (R-loop) via initial transcription of the ITR region. The transitions and transition rates in this system include: [1] the association rate of RNAP/σ recruitment to promoter DNA, transitioning the system from state 0 to 1 (r_1_); [2] the disassociation rate of RNAP/σ after it’s bound, transitioning the system from state 1 to 0 (r_-1_); [3] the rate of the closed-to-open conformational change, DNA melting, and stable R-loop formation, transitioning the system from state 1 to 2 (r_2_); [4] the rate of transcriptional bubble collapse and abortive initiation, transitioning the system from state 2 to 1 (r_-2_); and [5] the rate of productive transcriptional initiation, transitioning the system from state 2 to 0, whereby RNAP/σ transitions to the transcriptional elongation process, escapes from the promoter DNA, and frees up the promoter DNA to be available for binding to another copy of RNAP/σ (r_3_).

This coarse-grained representation of the transcription initiation process purposefully excludes several intermediate states and internal transitions. As discussed below, if there is no accumulation of internal states during transcriptional initiation (e.g. via significant pausing), then the addition of more internal states to the model is not beneficial and does not alter the free energy model formulation or coefficient values.

A chemical master equation (CME) can be formulated to describe the time-evolution of the probabilities of each state, considering the transition rates between each state. The CME for this system is:

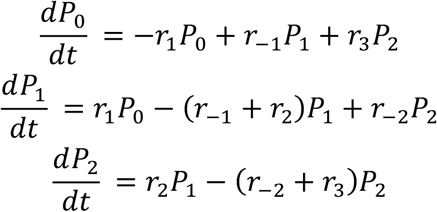

We also designate that the transcription initiation rate (TX_actual_) for a promoter variant will be the rate of promoter escape (r_3_) multiplied by the probability of state 2 (P_2_). Then, by assuming that the system has reached a “dynamic” steady-state where the probabilities of each state are not changing in time, we can set the derivatives in the CME to zero, solve for the probability of state 2 (P_2_) in terms of the transition rates and the probability of state 0 (P_0_), and then multiply by the transition rate r_3_ to yield the transcription initiation rate *TX*_*actual*_:

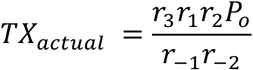

From transition state theory, we then recognize that the ratio containing the sequence-dependent transition rates can be reformulated in terms of a Gibbs free energy change between states 0 and 2, which we call ΔG_total_, as part of Boltzmann’s relationship. Here, Boltzmann’s constant β is a thermodynamic parameter that converts free energy differences into changes in state probabilities. In ideal (neutral, dilute) systems, β is 1/RT (R is the gas constant, T is temperature), but the value of β is not 1/RT in complex (charged, non-dilute) systems. We also recognize that our transcription rate measurements (*TX*_measured_) only provide a proportional measurement of the actual mRNA concentrations inside the system. We therefore create a proportionality constant K to denote this. We then observe that the steady-state probability of state 0 (P_0_) is indistinguishable from this proportionality constant. We therefore lump them together to yield the simple equation:

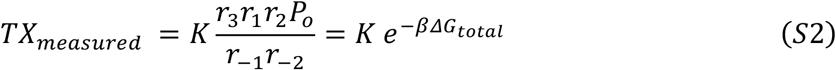

We then formulate a free energy model to calculate ΔG_total_ from sequence information. Our model considers each sequence-dependent interaction and then sums them together as linear, additive contributions. The free energy model is:

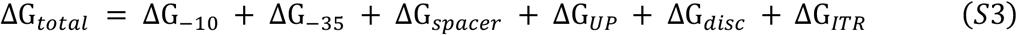

where ΔG_*-10*_ is the contribution from the -10 hexamer sequence, ΔG_*-35*_ is the contribution from the -35 hexamer sequence, ΔG_*spacer*_ is the contribution from the DNA spacer’s length and rigidity, ΔG_*disc*_ is the contribution from the discriminator sequence, ΔG_*ITR*_ is the contribution from the ITR sequence, and ΔG_*UP*_ is the contribution from the UP sequence.

We do not know the proportionality constant K, but by comparing the measured TX rates of the promoter variants to a designated reference promoter, we can remove the proportionality constant from the model training and prediction. To do this, we evaluate Equation S2 for a sample promoter variant (with measurement TX_measured_ and prediction ΔG_total_) and the reference promoter (with measurement TX_measured,ref_ and designated ΔG_total,ref_) and then subtract their left-hand-sides and right-hand-sides, yielding the equation:

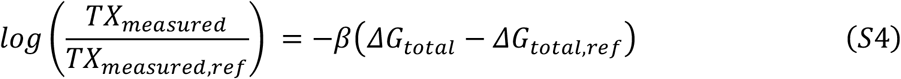

The choice of a reference promoter is somewhat arbitrary, but it should have a well-measured transcription rate. Here, we selected a reference promoter with a measured transcription rate closest to the log-mean-center of the dataset. The model itself is trained to predict the differences in ΔG_total_ between the promoter variants’ and the reference promoter (ΔG_total_ − ΔG_total,ref_), which leads to an interaction energy scale that spans both negative and positive values. In general, interaction energies have negative values when they are stronger than the average interaction strength. Interaction energies have positive values when they are weaker than the average interaction strength.

Next, we consider the assumptions made during model formulation and the experimental design aspects that ensure their validity. Assumption validity exists on a continuum and our objective here is to identify the best experimental conditions to maximize their validity. First, there is a pool of available RNAP/σ in the system that competitively binds to many DNA sites, including all promoter variants. This assumption is most valid when carrying out transcription measurements on many designed promoter variants (as in our study) or inside cellular conditions where many promoters co-exist, but would be least true if carrying out transcription rate measurements on a single promoter variant. In our *in vitro* transcription assay, the amounts of RNAP/σ and total DNA are also kept constant throughout the measurements. Second, the amount of RNAP/σ in the system is small enough, as compared to the amount of available DNA, that the promoter variants are not “saturated” and thus state 0 (the unbound state) is the predominant state. This assumption is most valid in our *in vitro* transcription assays and within typical *in vivo* (cellular) conditions, where the total amount of RNAP/σ is much smaller than the total amount of DNA. However, it would be least valid if excess amounts of RNAP/σ were added to the *in vitro* transcription assays. Third, we assume that transcriptional pausing does not occur in appreciable amounts across the promoter variants, which would lead to accumulation of state 1 and a reduction in state 0. The presence of pause sequences could make this assumption less true; however, we did not find any motif sequences in the discriminator or ITR regions whose inhibitory effect on TX rate could not be explained by other factors (e.g. R-loop thermodynamic stability). Fourth, the measured transcription rate of each promoter variant is proportional to the promoter escape rate (transition rate r_3_) multiplied by the probability of state 2 (stable promoter state), which is r_3_P_2_. We observe successful transcription when RNAP/σ transcribes each promoter variant’s unique barcode sequence, which is located in the 3’ UTR. By placing the barcode sequence in the 3’ UTR and not nearby the promoter, we remove the potential for barcode sequence-dependent changes in promoter escape that would make this assumption less true. We also utilize a short gene (*mRFP1*) to prevent appreciable amounts of RNAP/σ fall-off during transcription.

### Model Training using Machine Learning

Machine learning was used to train, test, and validate several linear models (Ridge Regression, LASSO, and ElasticNet) and parameterize the unknown interaction energies. To do this, we first enumerated a list of sequence motifs and biophysical characteristics (**Table S1**) that have the potential to contribute to the energetics of transcriptional initiation as quantified in our free energy model (**Equation S3**). There are 472 features in the exhaustive unpruned list. We divided our filtered dataset (the set of promoter variants with a single predominant TSS, see the Data Filtering section above) into a training set (4673 promoters, 90%) and an unseen test set (520 promoters, 10%). Transcription initiation rate measurements were log-transformed and converted to Z-scores, using the mean and standard deviation of the log-transformed transcription rates. We discuss the relationship between Z-score normalization and the biophysical modeling in a section below.

We then carried out model training and hyperparameter optimization (alpha, 1 parameter) using 10-fold cross-validation on the training set. The best models were then evaluated on the unseen test set. Accuracy metrics included the mean absolute error (MAE), mean squared error (MSE), and the squared Pearson correlation (R^2^). Feature importance was then carried out using one-feature drop analysis. Model training and testing was repeated several times, each time dropping individual features or sets of related features. The differences in model accuracy were calculated and features were pruned if they did not appreciably increase model accuracy. Feature drop analysis identified several features whose values were highly correlated with other features in the dataset, for example, AT content in the UP region and the minor groove width in the UP region. The final set of pruned features was identified by including all features that yielded appreciable improvements in model accuracy. When faced with equivalent feature set choices, we selected the smaller set of features with fewer unknown coefficients, which often relied on biophysical calculations. We also compared each model’s trained interaction energies against the canonical interactions known to have the most effect on transcription initiation rate, which we refer to as positive controls, and selected models that achieved 100% of these positive controls. Notably, the differences in interaction energies for the highest performing models were small, but the small differences could affect the rank-order of the top/bottom 10 sequence motifs, which affected their positive control successes. In **Table S2**, we show the accuracies of the top performing linear models using either the full or pruned feature sets. We selected the Ridge Regression model for use in this study, though optimized versions of each linear model type yielded high accuracies on the unseen test set.

The final set of pruned features included: (i) The energetic contributions from binding to the -35 and -10 hexamer were represented using one-hot encoding of the presence of each possible 3-nt motif within each hexamer. All possible 3-nt motif sequences (64 3-mers) are included and their presence or absence is a categorical feature, totaling 256 features. (ii) The energetic contributions from the spacer region were decomposed into two sets of features, including the spacer length (15 to 20 bp, each represented as a categorical feature) and the spacer’s DNA rigidity (a single numerical feature calculated based on the sequence-dependent persistence length of double-stranded DNA) ^26^. (iii) The energetic contributions from binding to the -10 extended motif were represented using one-hot encoding of the presence of each possible 2-nt motif located upstream of the -10 hexamer, totaling 16 categorical features. (iv) The energetic contributions from the discriminator region were represented using one-hot encoding of each possible 3-nt motif in the first 3 nucleotides of the discriminator region, totaling 64 categorical features. (v) The energetic contributions from the UP region were represented using the numerical value of the calculated minor groove width in the distal and proximal UP sites (each 10 bp long) ^23^. (vi) The energetic contributions from the ITR were represented by calculating the thermodynamic stability of the R-loop within the first 15 nucleotides of the ITR region (1 numerical feature), which is the free energy of the duplexed DNA-DNA complex subtracted by the free energy of the duplexed RNA-DNA complex. Numerical feature values are divided by the maximum possible value for normalization.

Additionally, during the initial model training, feature encoding relied on the positional identification of sequence motifs using the following procedure. The predominant transcriptional start site was determined from TSS Mapping measurements. The promoter variant sequence was then scanned, taking into account that discriminator lengths varied from 5 to 10 nt and spacer lengths varied from 15 to 20 nt. The positions of the -10 and -35 hexamer motifs were then identified by scanning across the sequence, evaluating a position weight matrix (PWM) tailored for each motif, and calculating a motif score. The positions with the highest motif score were labeled as the -10 or -35 hexamer. If two or more locations had equivalent motif scores, the motif location with the most optimal spacer length (17 nt) was selected. This procedure was tailored to function alongside our experimental promoter library design, which included the 4096 possible - 35 hexamer motif sequences alongside a consensus -10 hexamer as well as the 4096 possible -10 hexamer motif sequences alongside a consensus -35 hexamer. Even though these designs used a canonical spacer length of 17 nt, an alternate procedure that assumed that the spacer length was always a constant yielded scenarios where sequence changes led to shifts in the locations of the actual -10 or -35 motif and corresponding changes in motif sequence. Failing to account for these shifts led to mismatched interaction energies during model training, for example, non-canonical motifs having interaction energies similar to canonical motifs, created by the formation of a canonical motif with a shifted location.

### The Relationship between Biophysical Modeling and Z-score Normalization

To create our training and test datasets, we log-transformed the transcription rate measurements and applied Z-score normalization on the outcomes. Z-score normalization is a routinely used approach for improving the machine learning training process, whereby the outcomes in a dataset are transformed and re-scaled. The Z-score is:

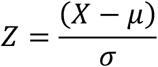

where X is the observed outcome, μ is the arithmetic mean of the observed outcomes, and σ is the standard deviation of the observed outcomes. Here, we also use a statistical thermodynamic model to predict the transcription initiation rate of a DNA sequence, according to:

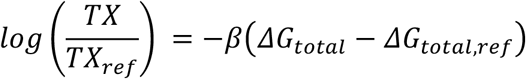

We show that these two relationships are directly related to one another, enabling us to determine the apparent thermodynamic parameter β, which is a model constant that converts free energies into state probabilities.

Specifically, we first log-transform the transcription rate measurements such that X is *log*(*TX*) for a particular promoter variant, μ is the mean of the log-transformed measurements, and σ is the standard deviation of the log-transformed measurements. Within the dataset, we selected the reference promoter such that its log-transformed measured transcription rate was very close to the mean of the log-transformed measurements (μ). Therefore, the X – μ in Z-score normalization is equivalent to 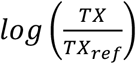 We then use this choice to equate the Z-score and biophysical model equations, such that

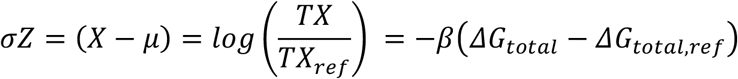

Here, we see that the standard deviation σ of the log-transformed measurements is equivalent to the negatively valued thermodynamic beta (−β) and that the Z-score used to train the model is equivalent to the apparent difference in free energies -*ΔG*_*total*_ − *ΔG*_*total*,.*ref*_. In other words, the standard Z-score transformation procedure for model training is equivalent to the use of state referencing in thermodynamic modeling of complex systems.

A key effect of the Z-score normalization procedure is to collapse the range of the actual and learned outcomes to aid in the model training process. This has the effect of collapsing the maximum magnitude of the interaction energies (**Figure 2**). However, we can revert the learned interaction energies back to their original scale by evaluating the biophysical model equation, which uses the learned thermodynamic beta. In **Table S5**, we show the apparent thermodynamic beta constants, learned for each dataset. Notably, the values are similar across all the *in vivo* datasets, but somewhat distinct for the *in vitro* dataset. There are physiochemical reasons for this difference, likely related to composition and molecular crowding differences between the *in vivo* and *in vitro* systems.

### Individual Promoter Construction and Isogenic Characterization

Individual promoters were constructed to test the effects of changing UP and ITR regions on transcription rates (**Figure 4A**) and to test the automated design of promoters using model predictions (**Figure 4E**). Promoters were constructed by synthesizing pairs of overlapping oligonucleotides and using PCR assembly to create insertion DNA cassettes. Cassettes and the mRFP1-expression plasmid pFTV1 were double-digested, purified via gel extraction, ligated, and transformed. Plasmid products were sequence verified using Sanger sequencing. Isogenic *E. coli* DH10B cells were transformed with each plasmid, following by quantification of their mRFP1 expression levels during exponential growth, using spectrophotometry and flow cytometry measurements. Cells were grown overnight in LB media supplemented with 50 ug/ml chloramphenicol, followed by a 1:100 dilution into 200 ul M9 minimal media supplemented with 50 ug/ml chloramphenicol within 96-well optical bottom microtiter plates. OD600 and mRFP1 fluorescence measurements were recorded every 10 minutes until cultures reached an OD600 of 0.20. Cultures were then serially diluted 1:10 into pre-warmed supplemented M9 media and grown again until they reached an OD600 of 0.20. 10 ul aliquots were then extracted and added to 190 ul PBS with 2mg/mL kanamycin to stop protein production. Flow cytometry was then carried out on the fixed cells to record their fluorescence distribution (BD LSR Fortessa). The mRFP1 fluorescence level was the arithmetic mean of the measured fluorescence distribution subtracted by autofluorescence, which was the arithmetic mean of the fluorescence distribution of wild-type *E. coli* DH10B cells. All designed promoter sequences, model calculations, and flow cytometry measurements are available in the **Supplementary Data**.

### Automated Promoter Sequence Design

We used an optimization algorithm (simulated annealing) to automatically design 16 promoter sequences with systematically varied transcription initiation rates, using the statistical thermodynamic model prediction within the optimization algorithm’s objective function. Designed promoter sequences were allowed to have variable TSS positions and use the surrounding upstream/downstream DNA sequence as part of the promoter. The surrounding upstream/downstream DNA sequences were kept constant for all promoter designs. The average promoter length was 77 bp and the average pairwise Hamming distance (the number of mismatched bp) across the 16 promoters was 57 bp, yielding an average sequence similarity of only 25.9%.

### Identification of Cryptic Promoters for Genetic Circuit Debugging

The statistical thermodynamic model was used to predict the transcription initiation rates across the 11-promoter genetic circuit (6793 bp), as studied and characterized in Borujeni et al. ^8^, and identify desired and undesired/cryptic transcriptional start sites (TSSs) on both its sense and antisense DNA strands. We labeled a location as a TSS when its predicted transcription initiation rate was at least 3-fold higher than the average transcription initiation rate per each strand. We used RNA-Seq measurements and RNAP flux inferences from Borujeni et al. ^8^ to assess the accuracy of these TSS predictions (**Figure 4H**). To do this, we counted the number of actual TSS peaks that were within 10 bp of the model-predicted TSSs and divided by the total number of model-predicted TSSs. This window threshold is reasonable considering that RNA-Seq measurements using 150 bp reads have limited positional precision and the RNAP flux inference calculation was shifted by up to 20 bp for known promoters with predominant TSSs.

## Supporting information

Supplementary Poster

## Author contributions

TL, AH and HMS conceived the study. TL designed and carried out the experiments. TL, AH, and HMS developed the algorithms and performed the data analysis. TL and HMS wrote the manuscript. All authors read, edited, and approved the manuscript.

## Data and materials availability

All promoter sequences, model calculations, and experimental measurements are available in **Supplementary Data**. A Python source code implementation of the statistical thermodynamic model of transcriptional initiation is available at https://github.com/hsalis/SalisLabCode. Next-generation sequencing read data files in fastQ format are available at NCBI with accession identifier PRJNA754118.

## Supplementary Information

**Figure S1:**
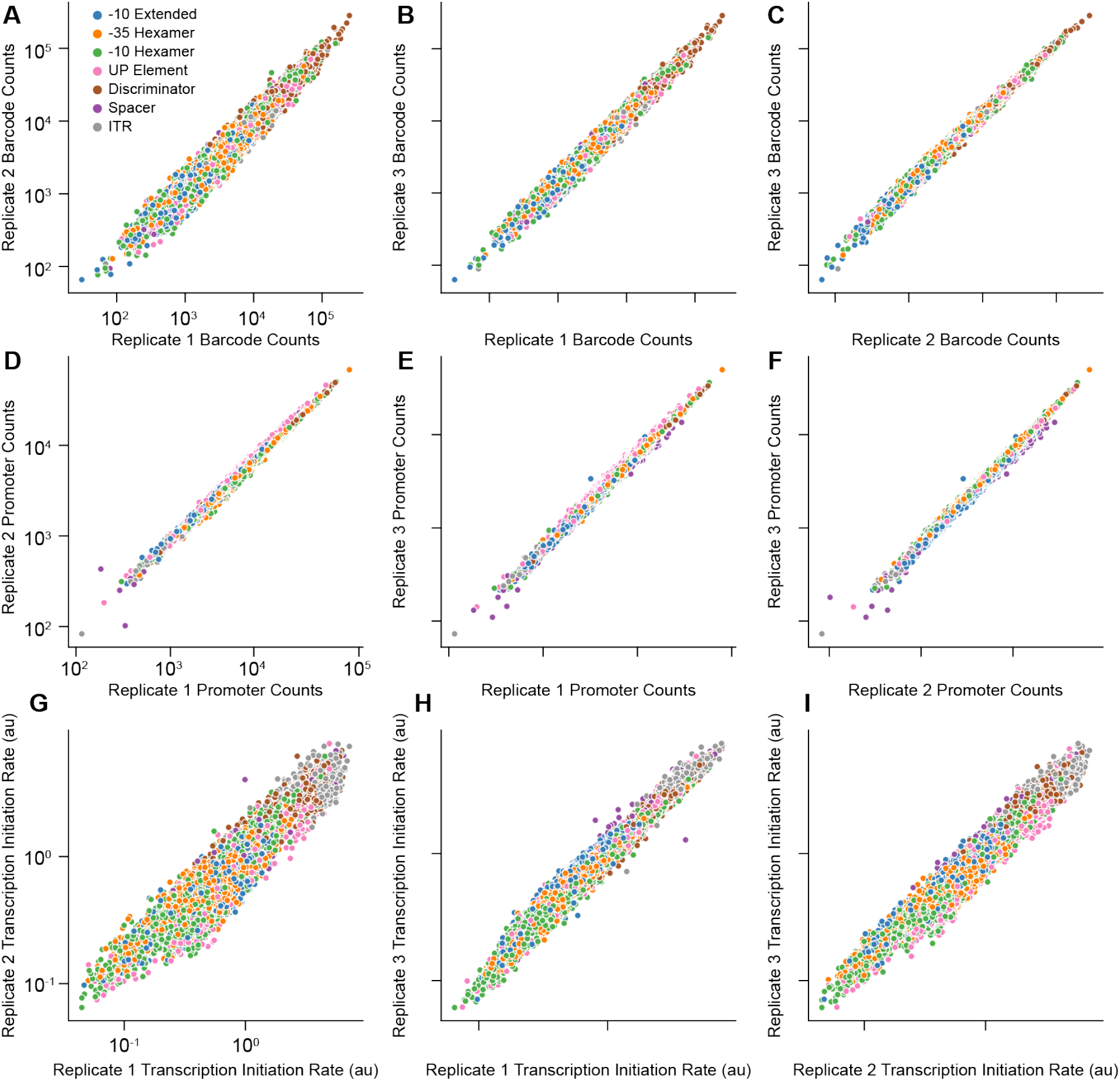
Replicate Correlations. RNA read count correlations for (**A**) replicate 1 vs replicate 2, (**B**) replicate 1 vs replicate 3 and (**C**) replicate 2 vs replicate 3 (*R*^2^ = 0.94, 0.97, and 0.99 respectively). DNA read count correlations for (**D**) replicate 1 vs replicate 2, (**E**) replicate 1 vs replicate 3 and (**F**) replicate 2 vs replicate 3 (*R*^2^ = 0.99, 0.99, and 0.99 respectively). Transcription Initiation Rate correlations for (**G**) replicate 1 vs replicate 2, (**H**) replicate 1 vs replicate 3 and (**I**) replicate 2 vs replicate 3 (*R*^2^ = 0.89, 0.94, and 0.97 respectively). Blue dots represent -10 extended promoter variants, orange dots represent -35 hexamer promoter variants, green dots represent -10 hexamer promoter variants, pink dots represent UP element promoter variants, brown dots represent discriminator promoter variants, purple dots represent spacer promoter variants, and grey dots represent ITR promoter variants.

**Figure S2:**
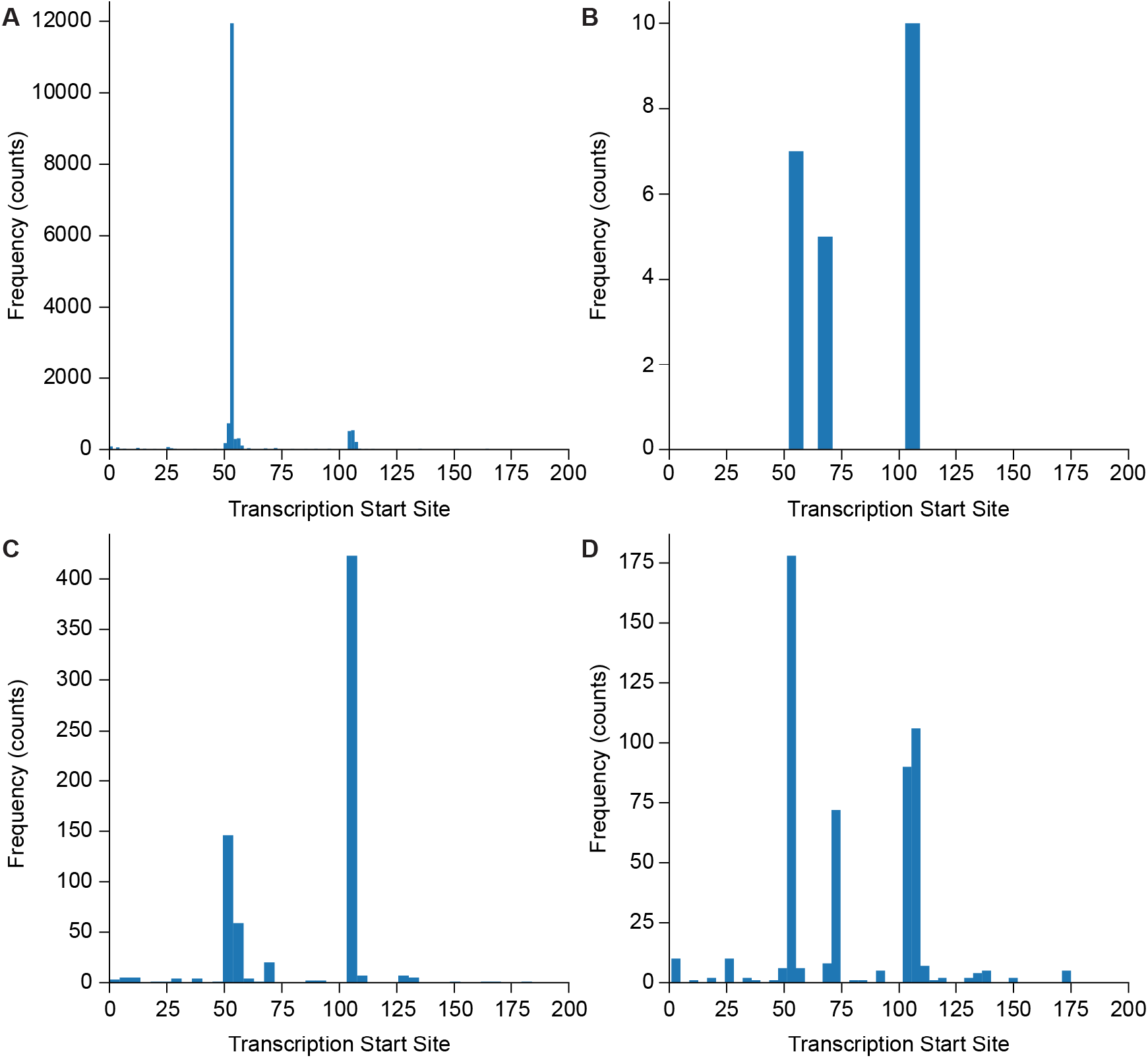
Transcription Start Site Profiles. (**A**), The measured TSS distribution of a variant that met all filtering criteria and exhibited a strong on-target peak. (**B**), The measured TSS distribution of a variant that did not meet the minimum read threshold. (**C**), The measured TSS distribution of a variant that had an off-target peak (position ∼105) with greater than twice the counts of the on-target peak (position 53). (**D**), The measured TSS distribution of a variant with multiple strong TSS peaks.

**Figure S3:**
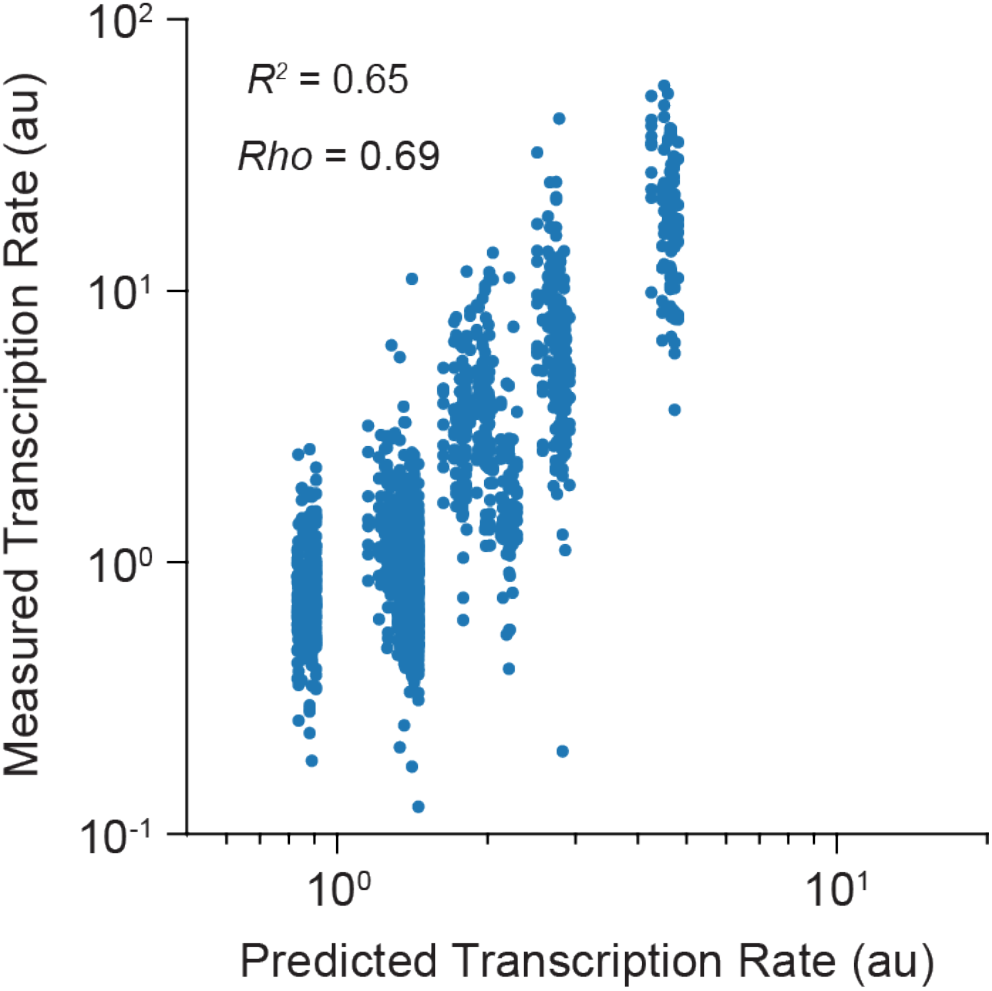
Model Accuracy on IPTG-inducible Promoters. Predicted vs. Measured transcription rates for 1493 genome-integrated promoters that contain lacO sites for IPTG induction, characterized *in vivo* by Yu et al. with 1 mM IPTG added ^29^.

**Figure S4:**
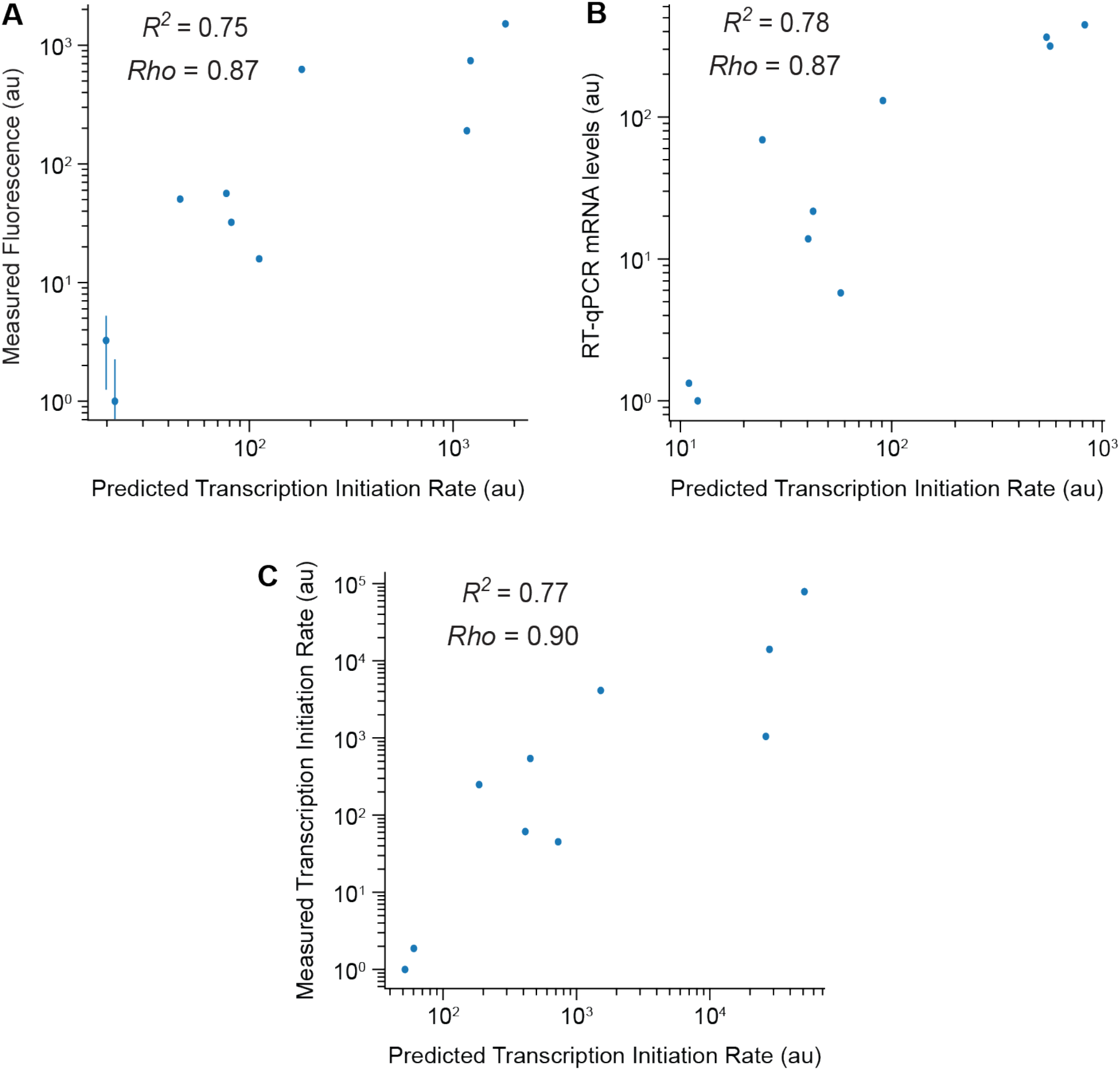
Model Accuracy on Selected Non-Repetitive Promoters with 3 Types of Measurements. (**A**), Predicted vs. Measured TX rates for the 10 promoters further characterized by Hossain et al. using flow cytometry. Error bars were created from duplicate measurements. (**B**), Predicted vs. Measured TX rates for the 10 promoters further characterized by Hossain et al. using RT-qPCR. (**C**), Predicted vs. Measured TX rates for the 10 promoters further characterized by Hossain et al. using read-based measurements.

**Figure S5:**
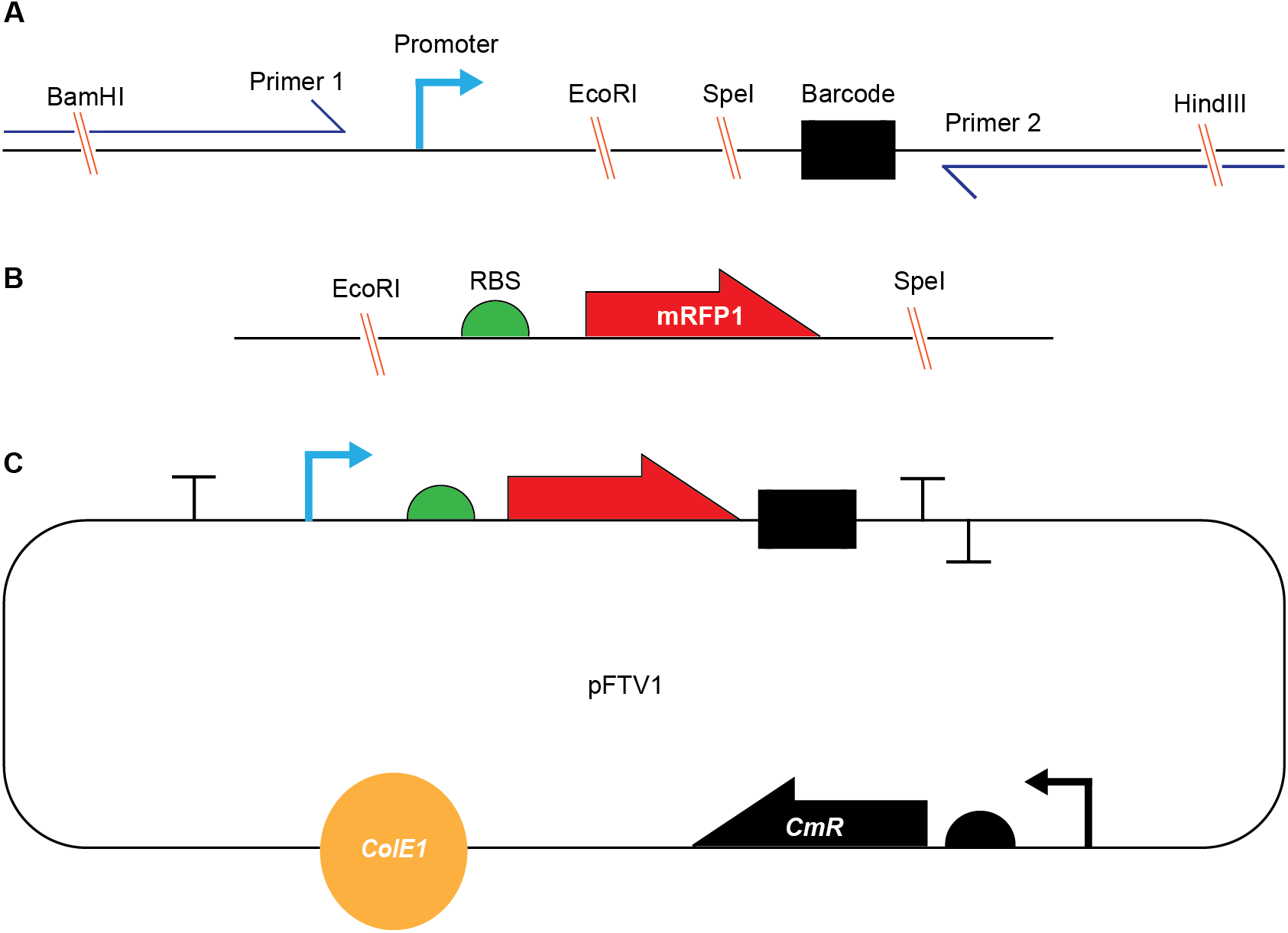
Designs and Plasmid Map. (**A**), Oligo Pool Library design with variable promoters. (**B**), Gene-block containing 5’ cleaving ribozyme, ribosome binding site and a reporter protein. (**C**), Final assembled plasmid used for promoter library characterization.

**Figure S6:**
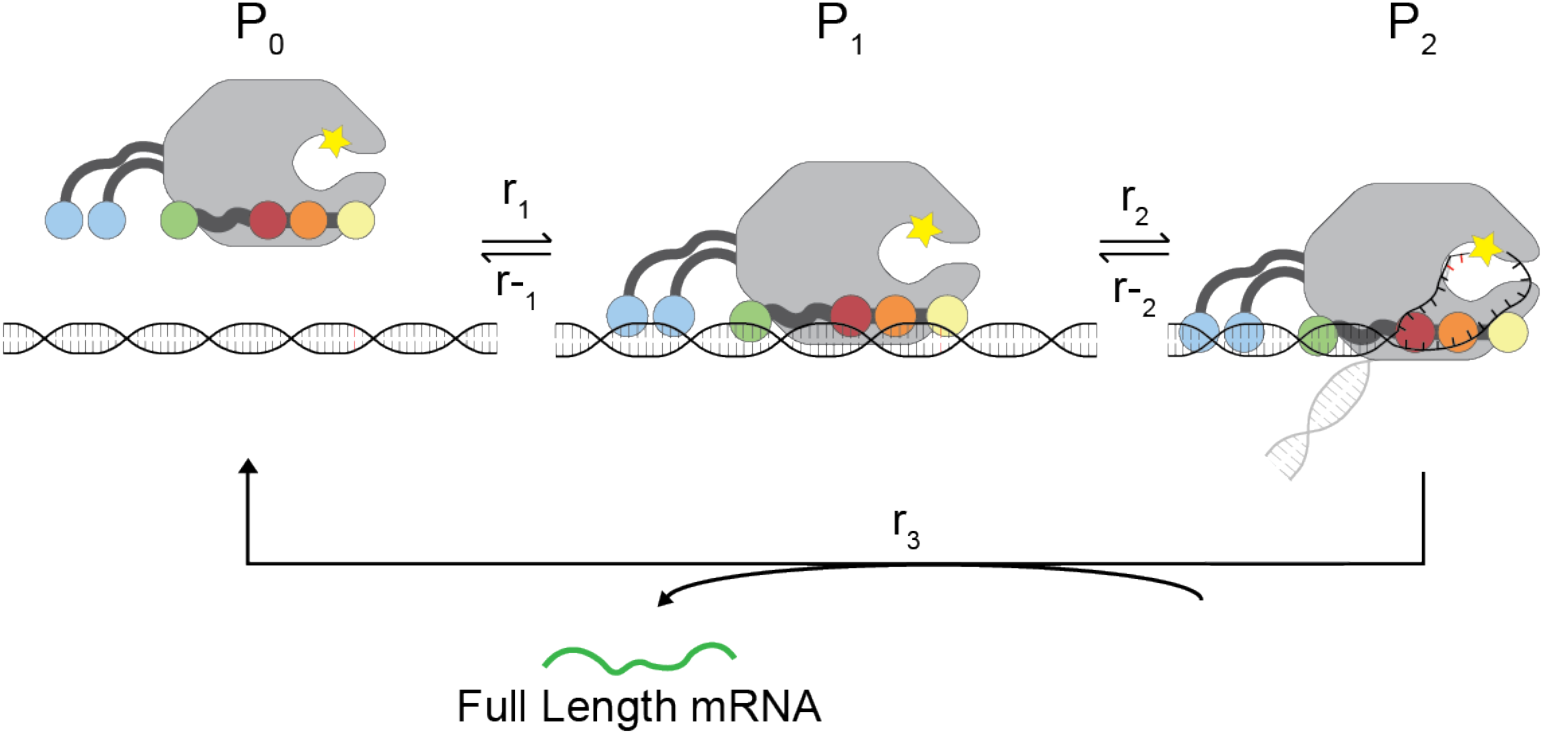
A 3-State Thermodynamic System. A coarse-grained representation of prokaryotic transcription initiation. P_0_ is the free RNAP and promoter DNA, P_1_ is the closed complex, and P_2_ is the open complex. Rates are denoted by r, where negative subscripts denote reverse rates and positive subscripts denote forward rates.

**Table S1:** Model Features

| **Feature Name, Feature Type** | **Number of Parameters** | **Number Retained<sup>a</sup>** |
| --- | --- | --- |
| -10 hexamer 3-mers | 128 | 128 |
| -35 hexamer 3-mers | 128 | 128 |
| Discriminator 3-mers | 128 | 64 |
| -10 extended 2-mers | 32 | 16 |
| Discriminator Length | 7 | 0 |
| Discriminator GC Content | 1 | 0 |
| Discriminator Purine Content | 1 | 0 |
| TSS 2-mers | 16 | 0 |
| Spacer Length | 6 | 6 |
| Spacer Stacking Free Energy | 1 | 0 |
| Spacer Torsional Energy | 1 | 0 |
| UP Groove Width (Distal and Proximal) | 2 | 2 |
| ITR Purine Content | 1 | 0 |
| ITR Pause-Element Location | 15 | 0 |
| UP A-tract Length (Distal and Proximal) | 2 | 0 |
| UP AT Content | 1 | 0 |
| DNA Rigidity | 1 | 1 |
| R-loop Strength | 1 | 1 |

<sup>a</sup> Retained parameters are the only ones present in the final model

**Table S2:** Model Selection

| **Model Type** | **# Variables** | **R2 Train** | **MAE Train** | **MSE Train** | **R2 Test** | **MAE Test** | **MSE Test** | **Correct / Total Positive Controls<sup>a</sup>** |
| --- | --- | --- | --- | --- | --- | --- | --- | --- |
| Ridge | 472 | 0.82 | 0.265 | 0.120 | 0.80 | 0.270 | 0.126 | 4/8 |
| Lasso | 472 | 0.82 | 0.264 | 0.120 | 0.80 | 0.270 | 0.126 | 4/8 |
| ElasticNet | 472 | 0.82 | 0.264 | 0.120 | 0.80 | 0.270 | 0.126 | 4/8 |
| Ridge | 346 | 0.80 | 0.274 | 0.133 | 0.80 | 0.281 | 0.131 | 8/8 |
| Lasso | 346 | 0.80 | 0.274 | 0.134 | 0.80 | 0.281 | 0.131 | 8/8 |
| ElasticNet | 346 | 0.80 | 0.274 | 0.133 | 0.80 | 0.281 | 0.131 | 8/8 |

<sup>a</sup> Positive controls were defined to match known consensus promoter sequence motifs <sup>12</sup>

**Table S3:**
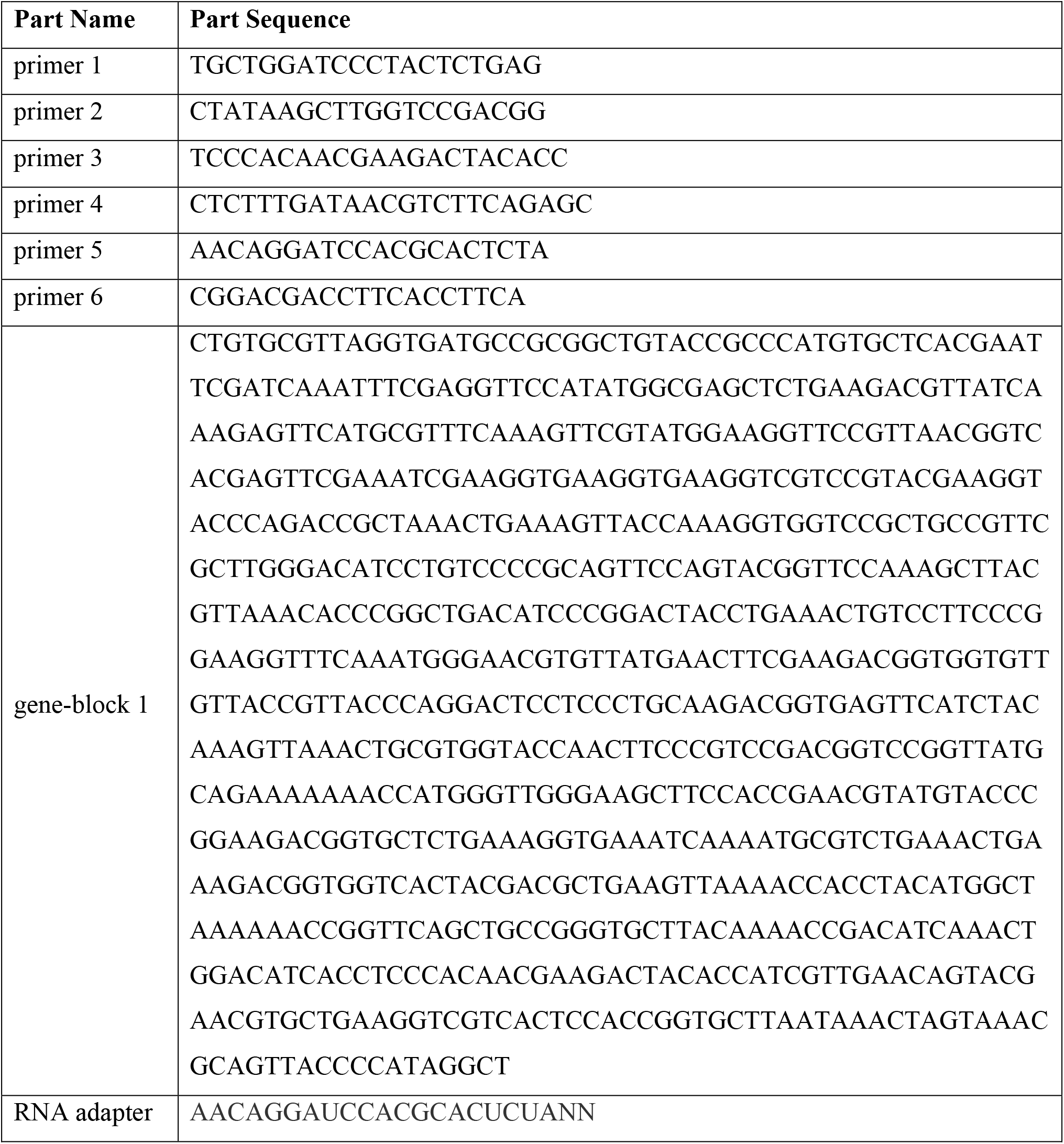
Primers, RNA Adapter, and DNA Fragment Sequences

**Table S4:** Model Performance Metrics

| **Data Set** | **Metric** | **Value** |
| --- | --- | --- |
| La Fleur et al. | Pearson Correlation | 0.79 |
| La Fleur et al. | Spearman Rank Order | 0.87 |
| La Fleur et al. | Mean Square Error | 0.23 |
| La Fleur et al. | Mean Absolute Error | 0.33 |
| Urtecho et al. | Pearson Correlation | 0.60 |
| Urtecho et al. | Spearman Rank Order | 0.78 |
| Urtecho et al. | Mean Square Error | 0.41 |
| Urtecho et al. | Mean Absolute Error | 0.54 |
| Hossain et al. | Pearson Correlation | 0.45 |
| Hossain et al. | Spearman Rank Order | 0.68 |
| Hossain et al. | Mean Square Error | 0.69 |
| Hossain et al. | Mean Absolute Error | 0.66 |
| Yu et al. | Pearson Correlation | 0.65 |
| Yu et al. | Spearman Rank Order | 0.69 |
| Yu et al. | Mean Square Error | 0.51 |
| Yu et al. | Mean Absolute Error | 0.55 |

**Table S5:** Apparent Values for Beta in Free Energy Model (Inverse Thermodynamic Temperature)

| **Data Set** | **Parameter** | **Value** |
| --- | --- | --- |
| La Fleur et al. | $\beta$ | 0.75 |
| Urtecho et al. | $\beta$ | 1.56 |
| Hossain et al. | $\beta$ | 1.86 |
| Yu et al. | $\beta$ | 1.07 |
| De Novo Designed | $\beta$ | 1.99 |
| Anderson et al. | $\beta$ | 1.82 |
| Hossain RT-qPCR (Figure S4B) | $\beta$ | 2.16 |
| Hossain Fluorescence (Figure S4A) | $\beta$ | 2.26 |
| Hossain Counts (Figure S4C) | $\beta$ | 3.45 |

