## Supplementary Poster for "Automated Model-Predictive Design of Synthetic Promoters to Control Transcriptional Profiles in Bacteria"

### The Promoter Calculator

#### a statistical thermodynamic model of transcription initiation in bacteria

Genetic System Sequence ⟶ Transcriptional Profile

##### List RNAP/σ Binding Sites

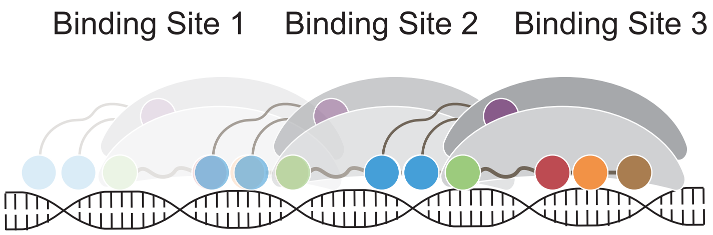


##### Calculate Total Free Energies

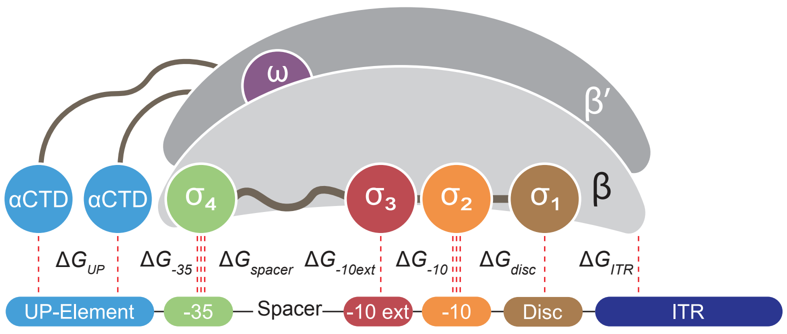


##### Predict Transcription Rates

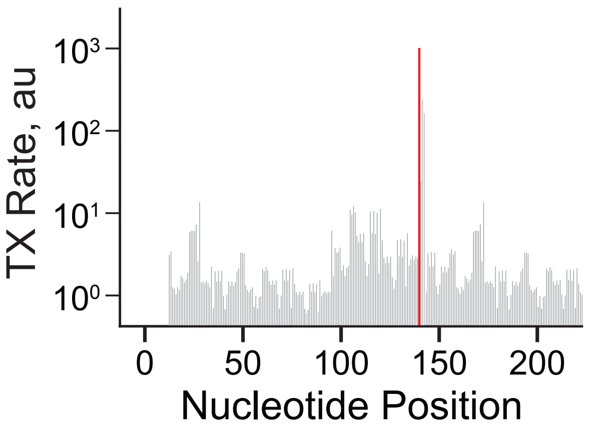


#### Free Energy Model for *Escherichia coli* RNAP/$\sigma^{70}$ Interactions with DNA

##### -35 Hexamer ($\Delta G_{-35}$)

| first 3 nt | | second 3 nt | |
|---|---|---|---|
| TTG | TGC | ACA | TGG |
| TTT | AAT | ACT | TTG |
| ATG | CCG | CCT | AAG |
| TAG | CGG | AAT | CGG |
| TGG | TAC | CCA | CTG |
| TTC | TCC | CAA | GAT |
| CTG | CAG | ATC | GAG |
| GTA | GAC | GTA | ATG |
| GTG | CTC | AAA | AAC |
| TCT | TCA | TAA | CCC |
| CTA | GGT | ACG | CGA |
| TAT | ACG | ATT | TTA |
| TTA | AGG | CTA | TGA |
| TCG | GCT | AGA | AGG |
| TGA | CAT | TGT | GTT |
| CTT | CCT | TTT | GTG |
| TAA | CAC | GAA | TCG |
| GTC | GAG | TCT | AGT |
| GTT | AAC | TCA | GGA |
| TGT | CGA | CTT | CTC |
| ATC | CCA | CAT | GGA |
| GCG | GGC | TAT | CGC |
| ATT | CAA | ATA | TAC |
| CGT | AAG | CAG | AGC |
| AAA | ATA | GCA | TTC |
| AGC | GGG | GCT | GCC |
| GGA | GCC | CCG | GGG |
| GAT | GAA | ACC | GTC |
| AGA | ACA | CAC | GAC |
| AGT | CCC | TAG | TCC |
| ACT | CGC | CGT | GGC |
| ACC | GCA | GCG | GGT |

##### -10 Hexamer ($\Delta G_{-10}$)

| first 3 nt | | second 3 nt | |
|---|---|---|---|
| TAT | ATC | AAT | AAG |
| TAA | ATG | TAT | CTA |
| TAC | AGC | ACT | TTA |
| TAG | GGT | AGT | AGA |
| AAT | TGC | CAT | TGA |
| CAT | TCC | ATT | GTA |
| GAT | AAC | GAT | CTC |
| GTA | TTG | TCT | AGC |
| CAA | TGT | GCT | GAA |
| TTA | TGG | TGT | ACG |
| CTA | CGA | ATA | TAC |
| CAC | TCT | CCT | CAC |
| ATT | TGA | GGT | GAC |
| GAC | CAG | TTT | CCG |
| CTG | AAA | GTT | GCC |
| GGC | GAA | CGT | GCG |
| CGG | AAG | CTT | GCA |
| CGC | GAG | ACA | TCG |
| GTC | ACT | AAA | CCC |
| GCC | CCA | AGG | GTC |
| GTG | GCA | TTC | CAG |
| GGG | TCA | AAC | TCC |
| CTC | AGT | ATG | GGA |
| GCG | GCT | TAG | TTG |
| ATG | CCC | CTG | TCA |
| ACC | GTT | GGG | GAG |
| ACG | TTC | GGG | GTG |
| CCG | TCG | CGA | CAA |
| AGA | TTT | TGG | GGC |
| GGA | CTT | ATC | CCA |
| AGG | CGT | ACC | TGC |
| ATA | CCT | TAA | CGC |

##### Discriminator ($\Delta G_{disc}$)

| first 3 nt | |
|---|---|
| GCT | GTG |
| TCT | TAG |
| GCA | AGG |
| CGC | GGC |
| CCT | AAT |
| CGA | ACA |
| CAC | CTC |
| TAT | TTC |
| TCC | AAC |
| TCG | TAC |
| CTG | GTC |
| GCG | CGG |
| ACC | TAA |
| TCA | GAT |
| CCG | GGG |
| CCC | CAA |
| ACT | TTA |
| GCC | AAG |
| AGT | CGT |
| CAT | TGA |
| TGC | CAG |
| ATA | AGA |
| CTA | GAA |
| TTG | ATC |
| CCA | AGC |
| AAA | TTT |
| GGA | GTA |
| ACG | TGG |
| TGT | GTT |
| GGT | GAG |
| GAC | ATG |
| ATT | CTT |

##### -10 Extended ($\Delta G_{-10ext}$)

**2 nt motif**

| | | | | | | | |
|---|---|---|---|---|---|---|---|
| TG | AT | CA | TA | TT | GT | AG | CC |
| GG | AA | CG | GA | AC | GC | TC | CT |

##### UP ($\Delta G_{UP}$)

| Groove Width Distal | | | | |
|---|---|---|---|---|
| 50 | 119 | 126 | 198 | 225 |

| Groove Width Proximal | | | | |
|---|---|---|---|---|
| 50 | 119 | 126 | 198 | 225 |

| AT Content | | |
|---|---|---|
| High | Med | Low |

##### ITR ($\Delta G_{ITR}$)

**RNA:DNA loop free energy**

-13.0 -9.0 -5.0

##### Spacer ($\Delta G_{spacer}$)

| Length | | | | | |
|---|---|---|---|---|---|
| 15 | 16 | 17 | 18 | 19 | 20 |

**Interaction Energy**: Strong -1.0, -0.5, 0.0, 0.5, 1.0 Weak

$$\Delta G_{total} = \Delta G_{UP} + \Delta G_{-35} + \Delta G_{spacer} + \Delta G_{-10ext} + \Delta G_{-10} + \Delta G_{disc} + \Delta G_{ITR}$$

$$TX = TX_{ref}\, exp(-\beta[\Delta G_{total} - \Delta G_{total,ref}])$$
